# CA3 circuit model compressing sequential information in theta oscillation and replay

**DOI:** 10.1101/2023.05.24.542204

**Authors:** Satoshi Kuroki, Kenji Mizuseki

**Author notes:** Contribution: Conceptualization, Investigation, Visualization, Writing - original draft, Writing – review and editing. Competing interests: No competing interests declared. Contribution: Supervision, Writing – review and editing, Funding acquisition.

## Abstract

The hippocampus compresses the sequential information in a theta oscillation cycle and short-term sequential activities during sharp-wave ripple while sleeping or resting; these processes are known as theta sequence and replay, respectively. The theta sequence is based on theta phase precession patterns of individual neurons. However, how these sequential neuronal activities are generated and how they store information about the outside environment remains unknown. We developed a hippocampal cornu ammonis (CA)3 computational model based on the biological CA3 circuit’s anatomical and electrophysiological evidence to address these. The model comprises theta rhythm inhibition, place input, and CA3-CA3 plastic recurrent connection. The model could compress the sequence of the external inputs and reproduce theta phase precession and replay, learn additional sequences, and reorganize previously learned sequences. A gradual increase in synaptic inputs, controlled by interactions between theta-paced inhibition and place inputs, explained the mechanism for sequence acquisition. This model highlights the crucial role of plasticity in the CA3 recurrent connection and theta oscillational dynamics, and hypothesizes how the CA3 circuit acquires, compresses, and replays sequential information.

## Introduction

The hippocampus contributes to episodic memory formation (Gordon, 1988; Scoville and Milner, 1957). Rodent spatial memory has been extensively studied as an animal model for episodic memory (Buzsáki and Moser, 2013; O’Keefe and Dostrovsky, 1971; O’keefe and Nadel, 1978; Tolman, 1948). In the hippocampus, place information is represented as a phase code within theta cycles of local field potential (LFP) through a phenomenon known as theta phase precession, where individual neurons firing timings within theta oscillations advance as the animal walks through the neurons’ place fields (O’Keefe and Recce, 1993; Skaggs et al., 1996). Consequently, sequential activity is manifested as the firing order within a theta cycle, forming theta sequences (Dragoi and Buzsáki, 2006; Foster and Wilson, 2007). The sequential activities are replayed during sharp-wave ripple (SWR) during sleep and quiet resting (Diba and Buzsáki, 2007; Foster and Wilson, 2006). The sequential activity is hypothesized to compress external world relationships in the order of seconds to 10 seconds into neural sequences in the order of 10 to 100 milliseconds. It enables more efficient processing, storage, and retrieval of information and integrates information into cognitive processes such as memory consolidation, abstraction, planning, and inference (Buzsáki, 2015; Drieu and Zugaro, 2019).

The hippocampus can be divided into several subregions, mainly the dentate gyrus (DG) and cornu ammonis 1 and 3 (CA1 and CA3), each of which plays an essential and distinct role in episodic and place memory. In particular, the CA3 is integral to generating sequential activity. The CA3 is a starting point for the ripple (Buzsáki, 1986; Buzsáki et al., 1983; Suzuki and Smith, 1988) and replay (Middleton and McHugh, 2016) activities. The CA3 receives input from the DG and projects it to the CA1. Input from the DG is very sparse in the CA3, however each input strength is large (Henze et al., 2002, 2000). There are more recurrent connections between the excitatory neurons in the CA3 than in the DG and CA1. Synapses between CA3 excitatory neurons are weaker, with a higher connection probability than synapses from DG to CA3 excitatory neurons. Both DG-CA3 and CA3-CA3 synapses show short- and long-term plasticity (Rebola et al., 2017). These anatomical features of CA3 circuits seem optimal for sequence learning (Banino et al., 2018; Jaeger and Haas, 2004; Laje and Buonomano, 2013; Sussillo and Abbott, 2009; Wang et al., 2018).

GABAergic neurons of the medial septum (MS) provide theta-paced inhibition to the hippocampal interneurons, which in turn control the firing timing of the excitatory cells (Freund and Antal, 1988; King et al., 1998; Petsche et al., 1962; Zutshi et al., 2018). Inhibitory cells also receive input from nearby excitatory cells and send them feedback inhibition (Rebola et al., 2017).

Several computational models for generating theta phase precessions (Chadwick et al., 2016; Geisler et al., 2010) and compressing sequences (Ecker et al., 2022; Jahnke et al., 2015; Nicola and Clopath, 2019; Reifenstein et al., 2021) have been proposed. However, no model has explained both these phenomena in a unified manner. Therefore, this study focused on the CA3 structure, which has many recurrent connections and is a source of SWRs, as a critical region to acquire sequence compression, and presents a mathematical model of how CA3 can memorize and replay the order of external place sequence.

## Results

We developed a minimal CA3 neural network based on anatomical evidence (Andersen et al., 2006; Rebola et al., 2017) to understand the fundamental mechanisms by which the CA3 recurrent circuit compresses sequential information (Figure 1A). Using a minimal model allowed for a detailed examination of the dynamics of individual neuronal units and their interactions (Chadwick et al., 2016; Izhikevich, 2010; Jensen et al., 1996; Reifenstein et al., 2021). The network consists of eight excitatory and one inhibitory neural unit. The excitatory neural units receive inputs from the DG, CA3 recurrent synapses, and the inhibitory unit. The DG input is sparse and large in a biological brain (Henze et al., 2002, 2000). The DG activity is also sparse, demarcating the input pattern more clearly, known as pattern separation (Yassa and Stark, 2011). To simplify the network, we assumed that each CA3 excitatory unit received only one DG input, although the synaptic weight was large (Figure 1A). Half of the DG inputs had place fields (order of units #1-2-3-4), and the others were silent (Figure 1B). The CA3 excitatory units were connected to each other, and the recurrent synapses were plastic. The CA3 excitatory units also received feedback inhibition from the CA3 inhibitory unit. The inhibitory neuron received excitatory inputs from CA3 excitatory units and theta-modulated inhibitory inputs from the MS (Freund and Antal, 1988; King et al., 1998; Petsche et al., 1962; Zutshi et al., 2018) (Figure 1A and C). We tested whether the network model can learn sequential activity within the network by experiencing the place order and whether it can replay the sequence without the external inputs.

**Figure 1.**
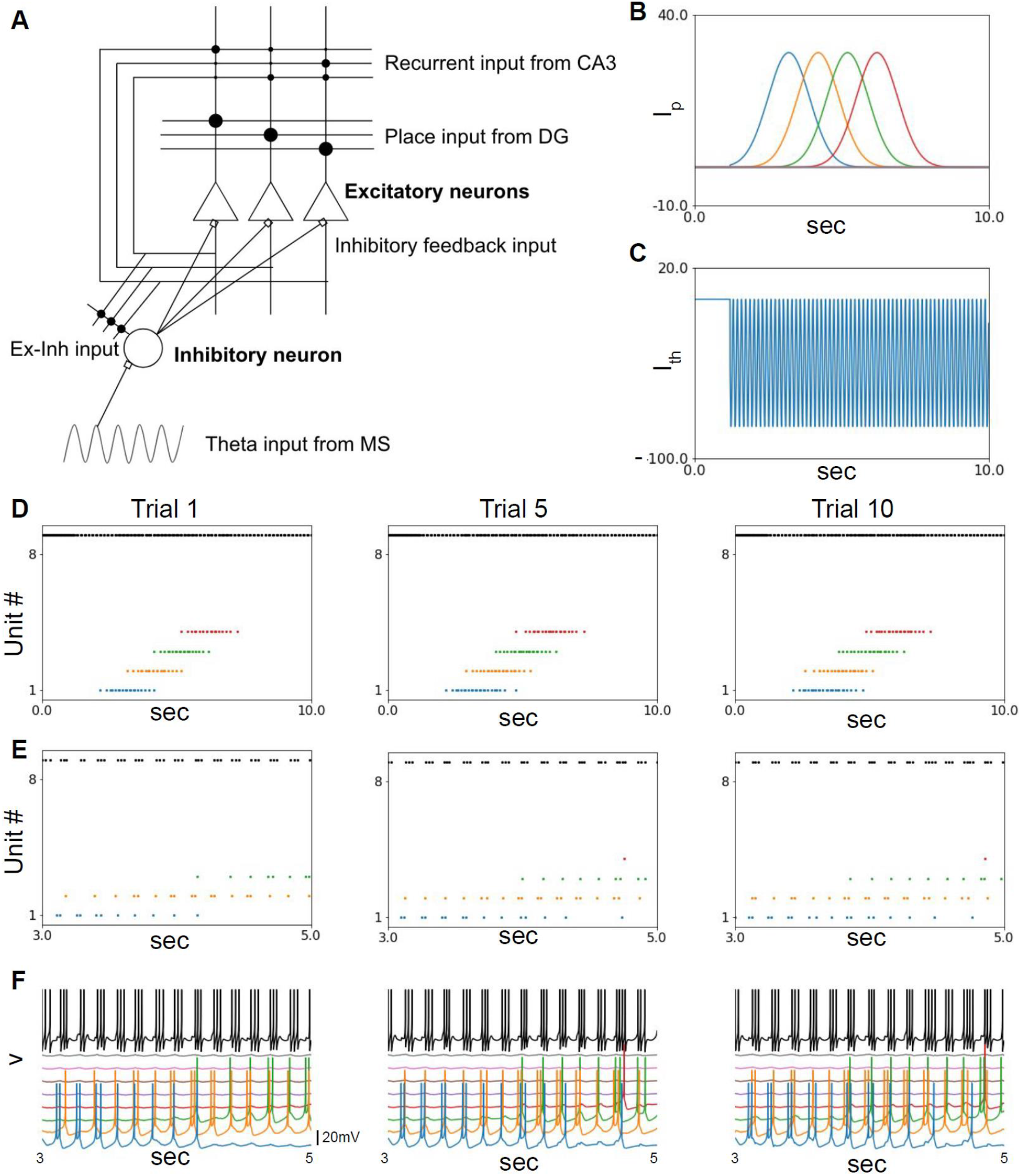
Small CA3 model (A) Schematic figure of model construction. The white triangle indicates the CA3 pyramidal neuronal unit, and the large white circle indicates the CA3 inhibitory neuronal unit. Lines indicate axons and dendrites. The small black and white circles on the lines are excitatory and inhibitory synapses, respectively. (B) Place inputs from the DG. (C) Theta input from the MS. (D) Spike raster plot during trials. The colored plots are spikes of the excitatory units, and the black plots are those of the inhibitory unit. (E) Zoom in of (D) from 3 to 5 s. (F) The membrane potential of each neural unit. The X-axis and color of each unit correspond to (D). CA, cornu ammonis; DG, dentate gyrus; MS, medial septum.

In this model, we set plasticity only in the CA3 recurrent synapses, not DG-CA3 synapses, to simplify the learning process. We selected the CA3 recurrent synapses because recurrent networks such as CA3-CA3 connections are suitable for learning sequential activities (Banino et al., 2018; Jaeger and Haas, 2004; Laje and Buonomano, 2013; Sussillo and Abbott, 2009; Wang et al., 2018). The plasticity rule was Hebbian. Activated synapses are tagged much longer than membrane potential changes (referred plasticity-related factor, see Material and methods) (Rogerson et al., 2014), which makes sequence learning with the Hebbian rule more efficient (Reifenstein et al., 2021). Furthermore, when inputs from the DG and another CA3 neuron arrive simultaneously, the neuronal response is much higher than the simple summation of the responses to these two inputs, known as superlinear coincident detection (Brandalise and Gerber, 2014; London and Häusser, 2005). We implemented these synaptic properties into the model (see Materials and methods). We also added the regularization term of the synaptic weight (maintaining summation of postsynaptic weights as the same constant value, see Materials and methods) to prevent overexcitation. The self-connection in the CA3-CA3 recurrent synapses was excluded.

The model was assumed to run on linear space with constant velocity for 10 trials (running session). During the running session, the plasticity on the CA3-CA3 recurrent synapse was changing the synaptic weights. Figures 1D–F show CA3 unit spikes (Figure 1D and E) and membrane potentials (Figure 1F) of both excitatory and inhibitory units in the first, middle, and last trials. Notably, as the trial progressed, the firing started earlier than in the previous trials, which is consistent with a previous experimental report (Mehta et al., 2000). The new spikes were generated because of the synaptic weight changes in the CA3-CA3 recurrent synapse.

The synaptic weights between neuronal units that received adjacent place inputs were increased with Hebbian plasticity, and those of the other synapses in the same postsynaptic unit decreased with regularization plasticity. The synaptic weights in the forward order (from earlier to later activated units) were more potentiated than those in the reverse order (from later to earlier activated units) (Figure 2A and B). The time course of synaptic weight potentiation differed between the order directions (Figure 2C). These plastic changes in synaptic weights accelerated the timing of spikes in Figure 1D–F.

**Figure 2.**
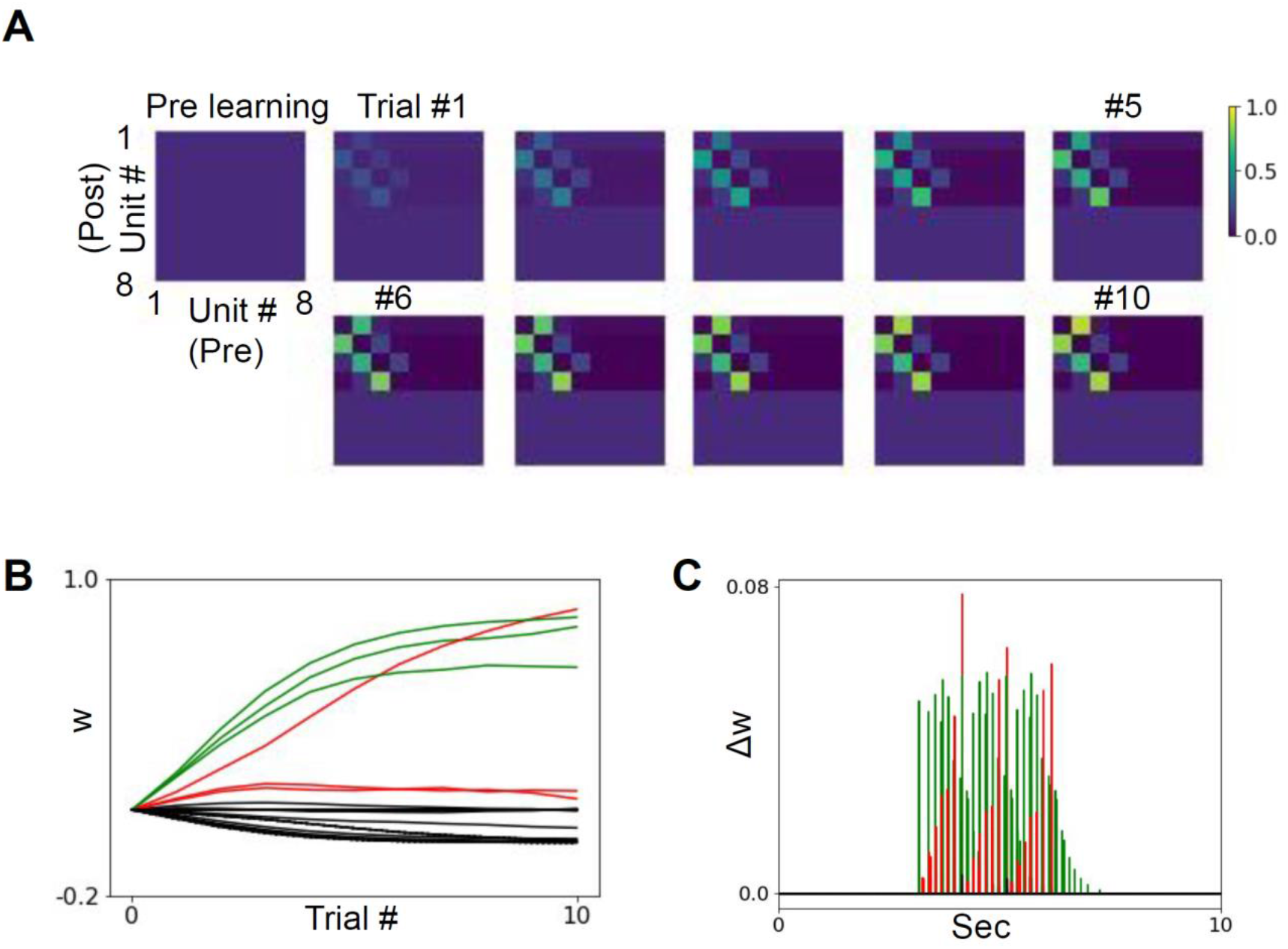
Synaptic weight change (A) Synapse weight of CA3 excitatory-CA3 excitatory recurrent connection in each trial. (B) The synapse weight as a function of trial numbers. The green lines are synapses in the forward order. The red lines are those in reverse order. The black lines are the other connections. (C) Synaptic weight changes (before regularization) during trial 1. The line color means the same as in (B). CA, cornu ammonis.

This model demonstrated theta oscillation as a population signal (Figure 3A and B) and theta phase precession-like firing pattern in the last trial (Figure 3D–F units #2–4) except the unit receiving first place filed inputs (Figure 3D blue dots, 3E and F unit #1). In contrast, the theta phase precession-like firing did not appear in the first trial (Figure 3C, E, and F). During the last trial, the excitatory units that received place inputs began firing at earlier positions in late phase than they did in the first trial, resulting in a negative slope of regression in units #2–4 (Figure 3C–F; two-tailed paired t-test, n=10; unit #1, p=0.005, t=3.667; unit #2, p=0.003, t=-4.083; unit #3, p=0.001, t=-4.704; unit #4, p=0.006, t=-3.529; Figure 3F). The enhanced synaptic weight of recurrent connections in Figure 2 caused these additional spikes. The model demonstrated the theta-precession pattern generation by learning the sequence.

**Figure 3.**
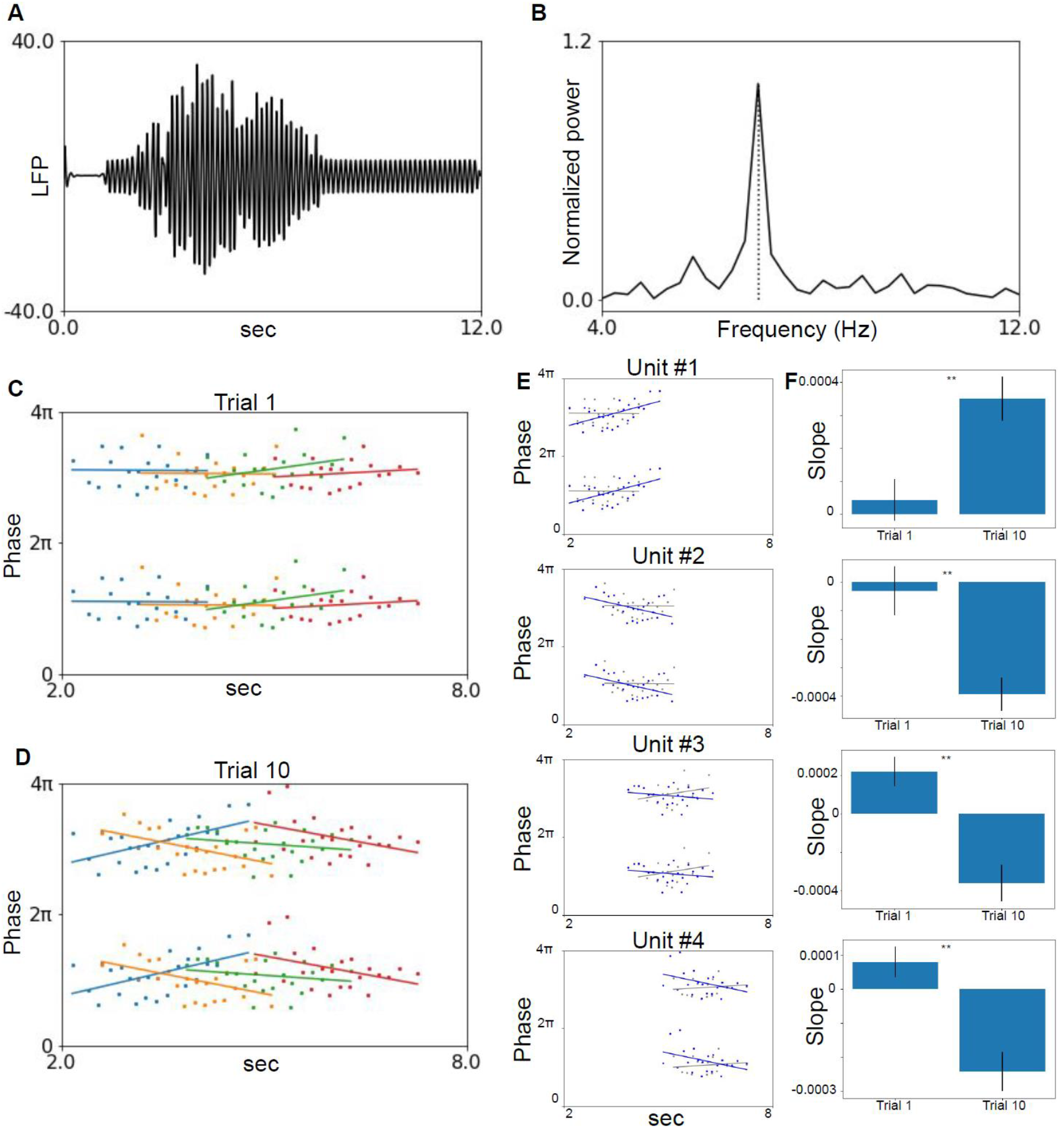
Phase precession (A) Local field potential (LFP): summation of membrane potentials of CA3 excitatory units trace. (B) Frequency spectrogram of the LFP. (C) Plot of theta phase and time (corresponding to the position) in trial 1. The colors of the points correspond to those in Figure 1D–F. (D) Same plot as (C) in trial 10. (E) Spiking phase vs. time plot in trials 1 (gray) and 10 (blue) of individual units are overlaid. (F) Slope values of trials 1 and 10 in individual units. The order of panels is the same as in (E). CA, cornu ammonis. ** p < 0.01

Next, we tested whether the model could replay the sequential activity without external place inputs. To simulate a resting state of animals, we added a noise input independently to each neural unit, cutting off the external place and theta inputs and shutting off plasticity in recurrent connections (resting session, see Materials and methods). After learning, the noise input occasionally induced random spikes, followed by sequential spikes from the associated units (Figure 4B). This did not appear before learning (Figure 4A). The lags between spikes among units that received place inputs (responded units) were smaller than those among other unit pairs after learning (Figure 4D) but not before learning (Figure 4C). By running multiple models for statistical testing, we confirmed that the densities of spike pairs within 100-ms lag were significantly higher among responded unit pairs (two-tailed paired t-test, n=10, p<0.001, t=20.4, Figure 4E upper) but not among other types of pairs (p=0.98, t=0.024, Figure 4E middle; p=0.35, t=0.97, Figure 4E lower). The difference in peak positions of place inputs during learning and lags during replay after learning between the responded units were significantly positively correlated in -200 to 200 ms lag (Figure 4F right, linear regression: n=104, slope=47.42, intercept=4.43, r=0.44, p<0.001), indicating this model mainly exhibited forward replay. We excluded unit #1 from the regression analysis because it showed different patterns in theta phase precession compared with other units. Emergency of the replay was also due to strengthening of synaptic weights between neighbor place field units (Figure 2). The model successfully reproduced replay spike sequences after learning the place sequence.

**Figure 4.**
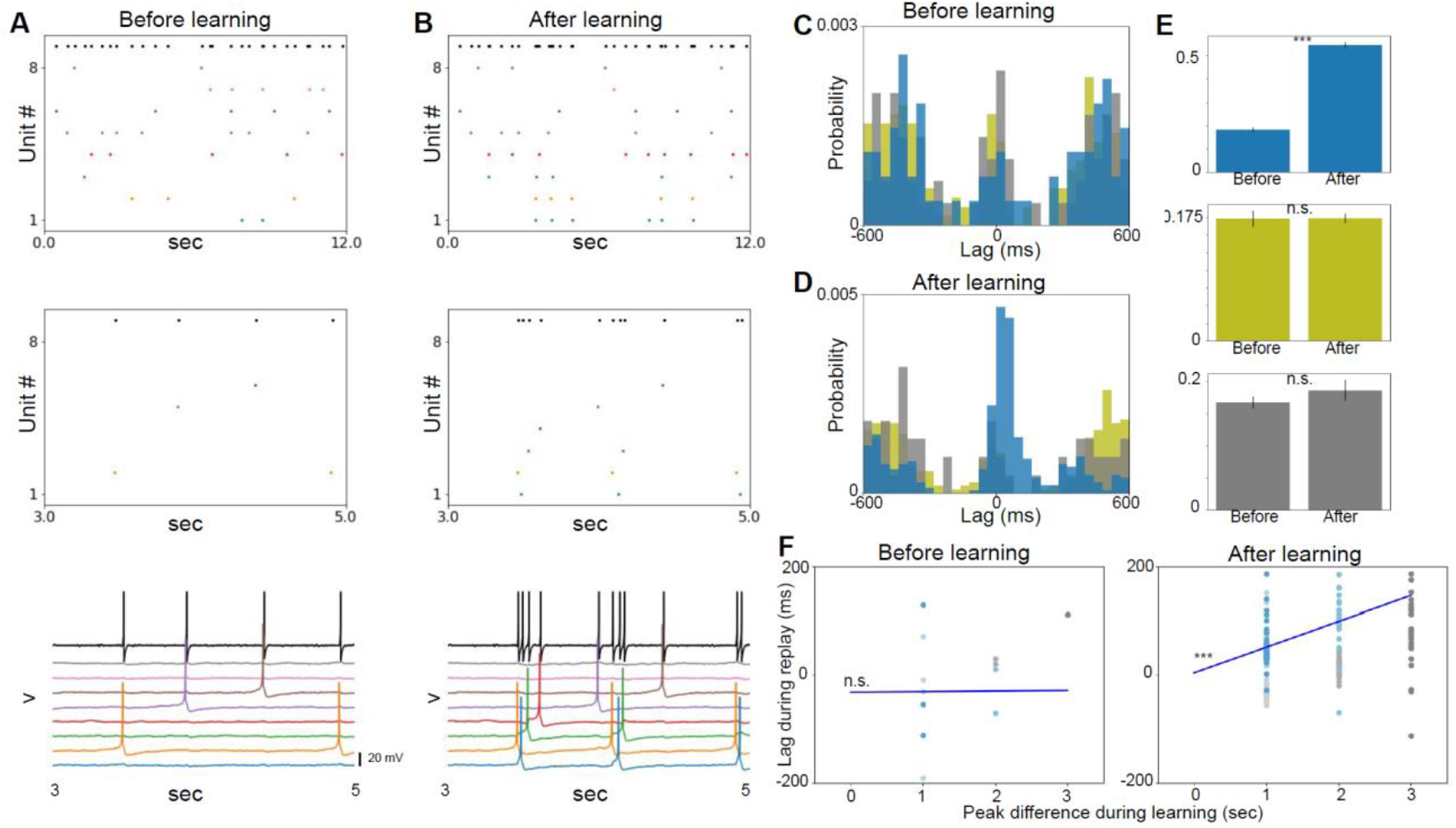
Replay (with 4 supplements) (A) Top: spike plot in the resting session with noise input before learning. The colored dot indicates the excitatory units, and the black dot is the inhibitory unit. Middle: zoom in of the upper panel for 2 s. Bottom: membrane potentials of each neural unit. The x-axis and color of each unit correspond to the middle panel. (B) is the same as (A) in the resting session after learning. (C) Crosscorrelogram of CA3 excitatory units of resting session before learning. (D) Crosscorrelogram of the resting session after learning. Blue shows pairs among responded (they receive I_p_) units during the running session. Yellow shows the pairs between responded and non-responded units. Gray shows pairs among non-responded units. (E) The densities of spike pairs within 100-ms lag among responded unit pairs (upper), between responded and non-responded pairs (middle), and among non-responded pairs (lower). (F) Scatter plot of the difference in peak positions of place inputs during learning and lags during replay before (left) and after (right) learning between responded units. The blue points show pairs excluding unit #1, and the gray points show those including unit #1. The blue lines show the linear regression results of the blue points. CA, cornu ammonis. *** p < 0.001, n.s. p > 0.05

We tested the models by partially altering the elements of the original model (see Materials and methods). A model with self-connection showed theta phase precession but not the sequential replay like in the original model (Figure 4—figure supplement 1). A model without coincident detection did not exhibit the theta phase precession or replays (Figure 4—figure supplement 2). A model with a short time constant of a plasticity-related factor did not show theta phase precession and replays (Figure 4—figure supplement 3). A model without the theta input from MS displayed the replay but not the theta phase precession (Figure 4—figure supplement 4). The weakening synaptic weight caused deficits in the model without coincident detection and in the model with a short plasticity-related factor time constant. By increasing the learning rate (10 times larger), the models recovered the replay in a forward or reverse order to some extent (Lower panels in Figure 4—figure supplements 2 and 3). These elements are necessary to acquire the theta phase precession and replay efficiently.

We tested the ability of the model to learn an additional sequence (order of units #4-5-6-7, Figure 5A), following the initial sequence learning (Figures 1–4). The first DG input of the new sequence was the same as the last one of the initial sequence set (unit #4, Figure 1B red line), while the other DG inputs differed from those in the initial sequence set. Similar to the initial sequence learning, the model demonstrated a shift in spike timing across trials in the additional learning stage (Figure 5B). The weights of the pairs within the additional sequence were strengthened, while the weight strengths within the initial sequence were maintained to some extent (Figure 5C). In addition, the model exhibited theta phase precession in the units that had learned the additional sequence (Figure 5D). During the resting session, the model displayed replay-like activity of the additional sequences set mainly in a forward order (slope=54.2, intercept=-10.7, r=0.43, p<0.001, n=122) and that of the initial sequence set with losing the direction of the order (slope=-36.8, intercept=70.3, r=-0.26, p=0.11, n=40) (Figure 5E–G). These results suggest that the model can learn additional sequences while retaining the associative information of the initial sequence.

**Figure 5.**
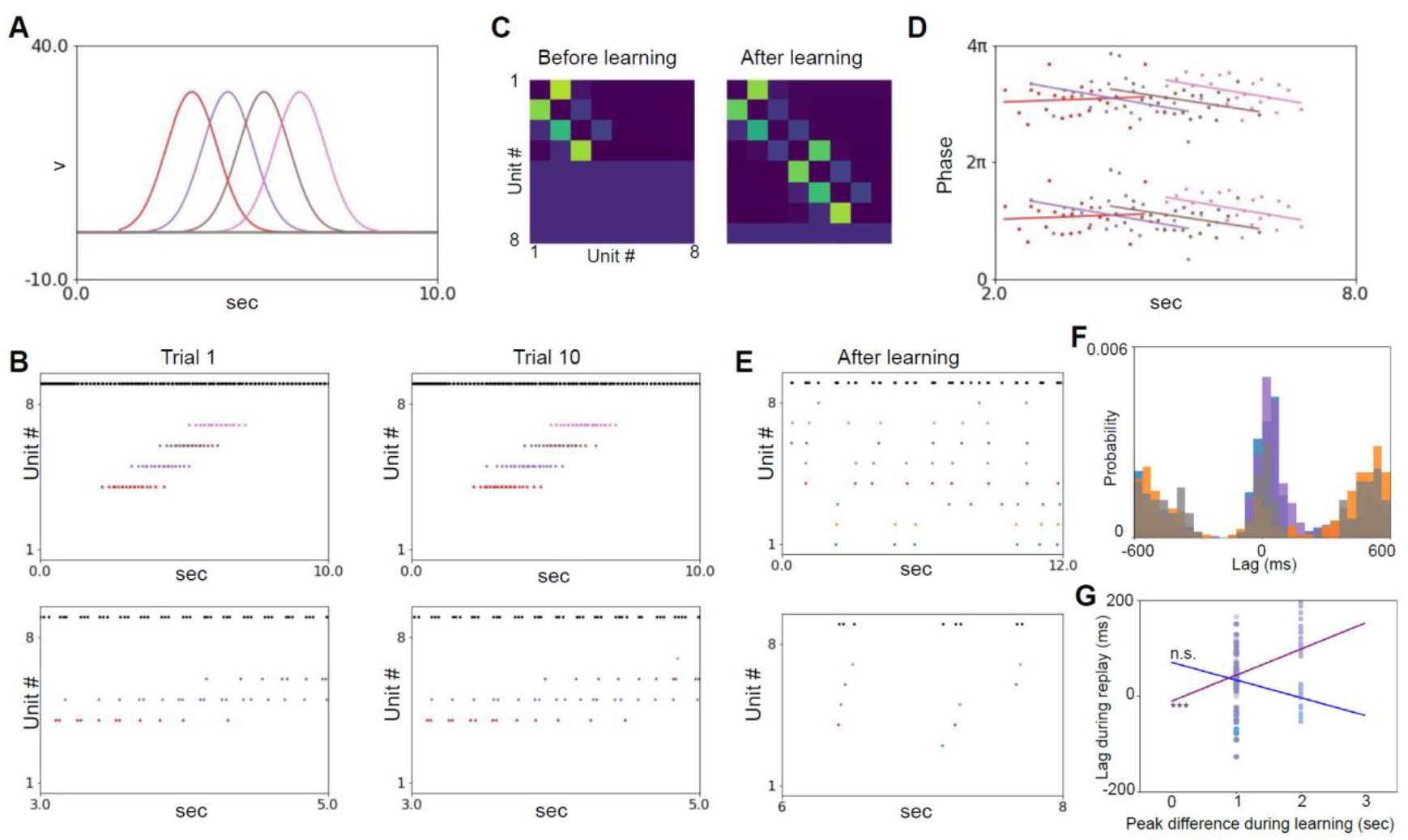
Additional learning (A) Place inputs from the dentate gyrus. (B) Upper: spike plot of trials 1 and 10. Bottom: zoom in for 3 to 5 s of the upper panels. The colored dots show excitatory spikes, and the black dots show inhibitory spikes. The colors correspond to those in Figures 1–4. (C) Synapse weights before (left) and after (right) the additional learning. (D) Phase precession after learning. (E) Spike plot of replay in the resting session with noise input after learning. The lower panel is zoomed in of the upper panel for 2 s. (F) Crosscorrelogram of the resting session after learning. Purple shows pairs of additional sequences. Blue shows pairs of the initial sequence (Figure 1– 4). Orange shows pairs between the initial and additional sequences. Gray shows the other pairs. (G) Scatter plot of the difference in peak positions of place inputs during learning and lags during replay after the additional learning. The purple points show pairs of the additional sequence (excluding unit #4), and the blue points show pairs of the first sequence (excluding unit #1). The purple and blue lines show the linear regression results of each colored point. *** p < 0.001. n.s. p > 0.05.

The ability of the model to relearn the new sequence, which was a mixture of the previous and new place fields, was also examined (order of units #8-1-3-6, Figure 6A) after the initial sequence learning (Figures 1–4). The second- and third-place inputs in the relearning sequence set were the same as the first-(unit #1) and third (unit #3)-place inputs in the initial sequence set. During the earlier trial, the units of the initial sequence set were spiked (yellow and red dots in Figure 6B left), but these activities disappeared after relearning (Figure 6B right). The synaptic weights underwent reorganization to accommodate the new relearning sequence (Figure 6C). The connections between the initial sequence set remained, although the synaptic weights were weakened. The theta phase precession of the relearning sequence was observed (Figure 6D). During the resting session, the model showed replay-like activity of the relearning sequence set in the forward order (slope=72.1, intercept=-27.6, r=0.63, p<0.001, n=80) and that of the initial sequence set with losing the direction of the order (slope=22.1, intercept=-0.54, r=0.17, p=0.114, n=86) (Figure 6E–G). These results indicated that the model can adaptively reorganize new sequence sets.

**Figure 6.**
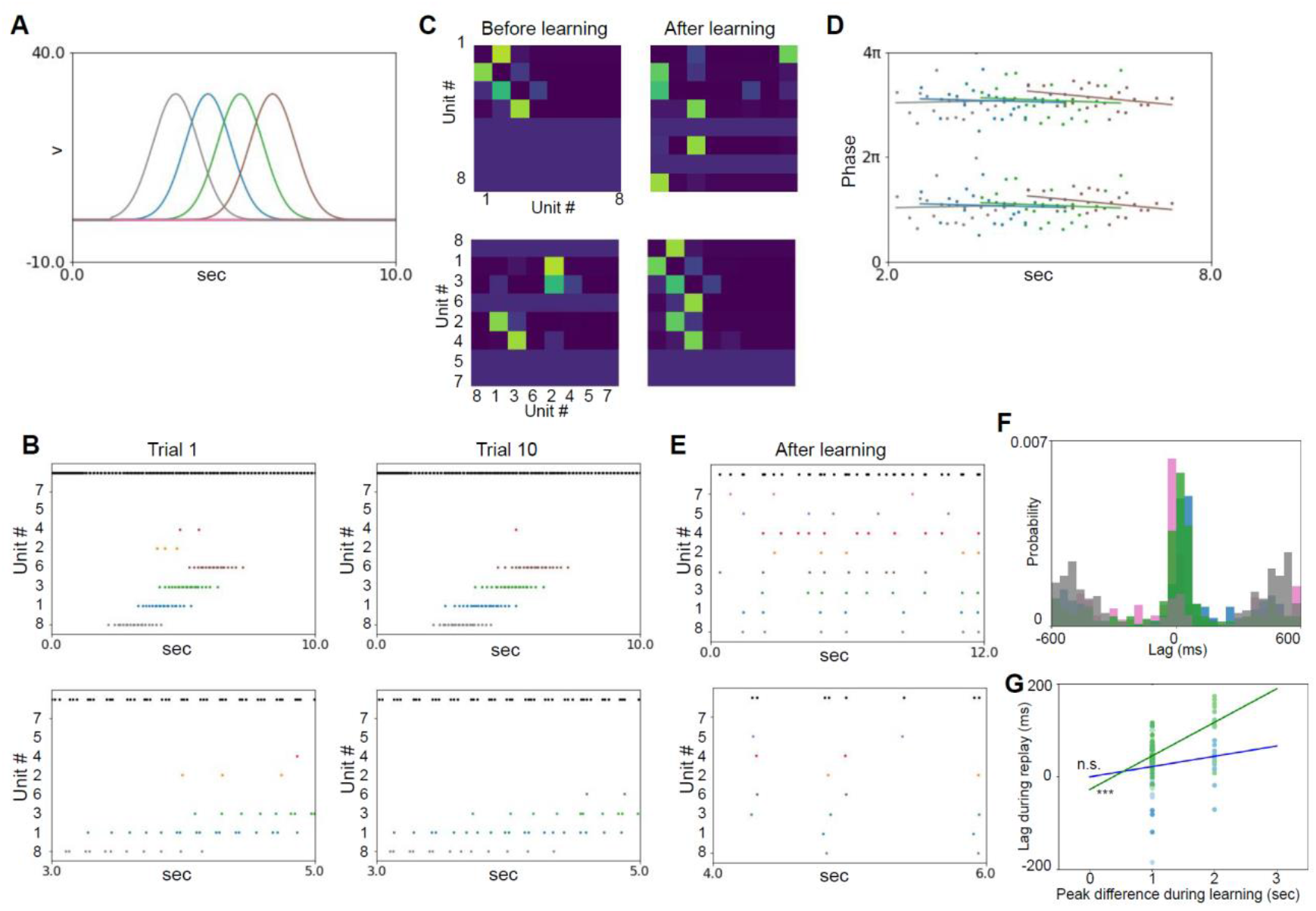
Relearning (A) Place inputs from dentate gyrus. (B) Upper: spike plot of trials 1 and 10. Bottom: zoom in for 3 to 5 s of the upper panels. The colored dots show excitatory spikes, and the black dots show inhibitory spikes. The colors correspond to those in Figure 1–4. (C) Synapse weights before (left) and after (right) the relearning. The upper panels are in the original order, and the lower panels are in the order of responded units in the relearning. (D) Phase precession after relearning. (E) Spike plot of replay in resting session with noise input after learning. (F) Crosscorrelogram of the resting session after relearning. Green shows pairs of relearning sequences. Blue shows pairs of the initial sequence (Figures 1–4). Magenta shows pairs between the initial and the relearning sequences. Gray shows the other pairs. (G) Scatter plot of the difference in peak positions of place inputs during learning and lags during replay after the relearning. The green points show pairs of the relearning sequence (excluding unit #8), and the blue points show pairs of the initial learning sequence (excluding unit #1). The green and blue lines show the linear regression results of each colored point. *** p < 0.001. n.s. p > 0.05.

We tested the minimal network model, in which synaptic connections were predetermined, in the above study. Notably, many more neurons are involved, and synaptic connections are probabilistic in the actual CA3 circuit. To replicate this in a more realistic simulation, we constructed a large model using the Brian 2 simulator (Stimberg et al., 2019). Similar to the small network, this model received two types of external input: the place input from DG to the CA3 excitatory units (Figure 7A) and theta inhibitory input from MS to the CA3 inhibitory units (Figure 7B). A tiny population of CA3 excitatory units was connected with and spiked by the DG place inputs (the probability of the DG-CA3 excitatory connection *p*_*pe*_ = 0.005), while all CA3 inhibitory units showed theta-locked burst spikes (Figure 7C). The CA3-CA3 recurrent synaptic connections were also probabilistic (the probability of the recurrent connection *p*_*rec*_ = 0.1) but much denser than the DG-CA3 synapses, and plasticity was implemented using the Hebbian rule. We selected the responded excitatory units (referred to as responded units) to observe the formation of the sequential activity (Figure 7D). Some units exhibited an earlier onset of activity during the last trial due to the learning (Figure 7E), and some synaptic weights between the responded units were enhanced (Figure 7F). Theta phase precession was also observed in the units after learning (Figure 7G and H). The model demonstrated replay activity during the resting session (Figure 7J–L). The results produced by the large network model were consistent with the results of the small network model.

**Figure 7.**
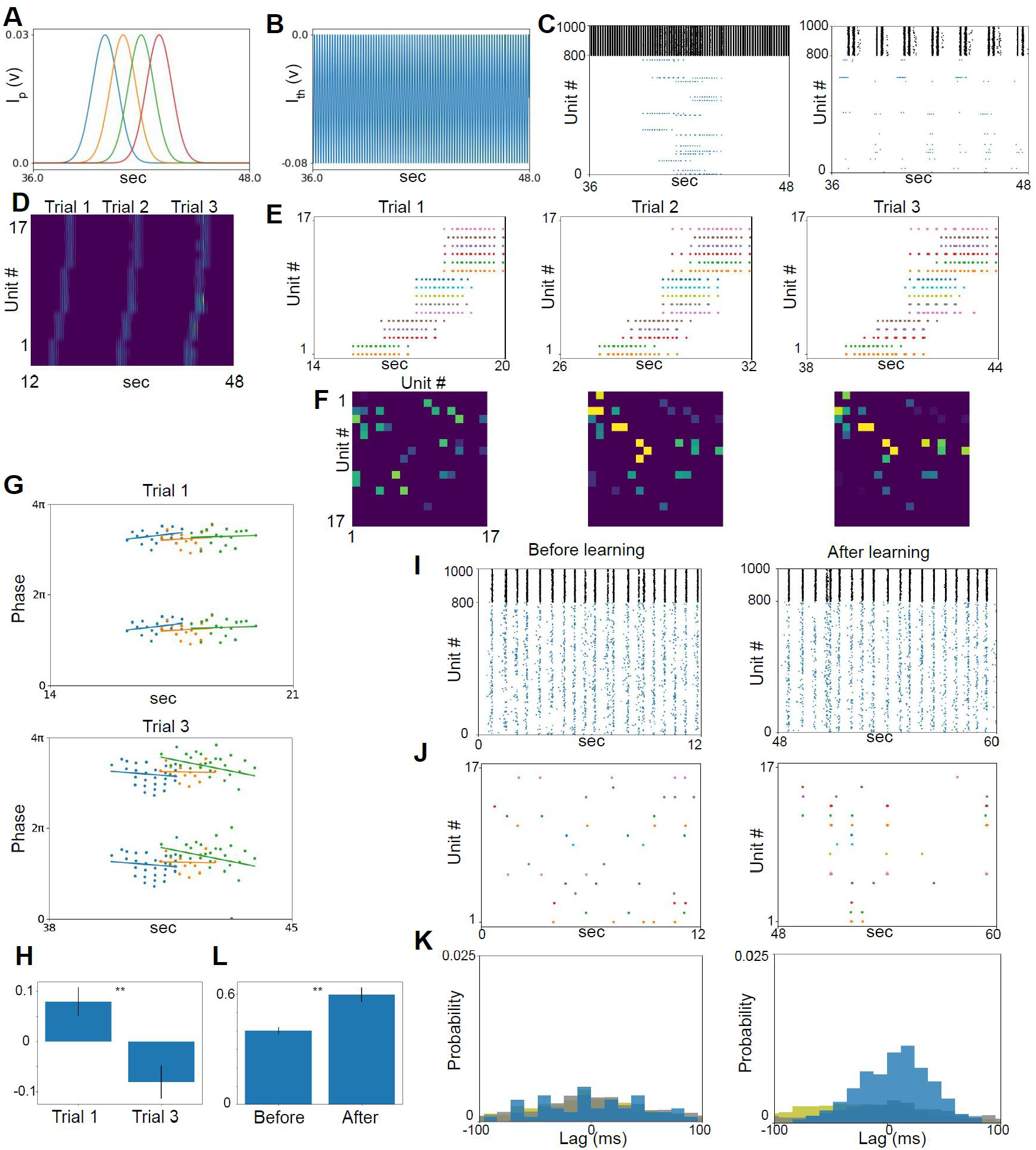
Large-scale network (A) Place input from the dentate gyrus. (B) Theta input from the medial septum. (C) Spike raster plot of excitatory (blue) and inhibitory (black) units. (D) The color plot of the spike rate of excitatory units responded to place inputs. They are sorted by peak response position and concatenated from trials 1 to 3. (E) Spike raster plots of the responded units from trial 1 (left) to 3 (right). Notably, the first spikes of some units proceeded in later trials. (F) Synaptic weights between the responded units from trial 1 (left) to 3 (right). (G) Theta phase precession in trials 1 (upper) and 3 (lower). (H) Slope values of responded units in trials 1 and 3 with multiple models (paired t-test, n=57, t=-3.535, p=0.001). (I) Spike raster plot of all units in resting session with noise input before (left) and after (right) learning. The color is the same as in (C). (J) Spike raster plot during the resting session of responded excitatory units. (K) Crosscorrelogram in the resting session. Blue shows unit pairs between responded units. Yellow shows unit pairs between responded and non-responded units. Gray shows unit pairs between non-responded units. (L) Densities of spike pairs within 100-ms lag among responded unit pairs before and after learning with multiple models (paired t-test, n=15, t=4.283, p=0.001). ** p < 0.001

Next, we analyzed the dynamics of each unit type when the external inputs 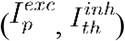 were supplied. We referred to the parameters of the above models’ excitatory and inhibitory neural units (Izhikevich, 2010). The dynamics of the Izhikevich model can be depicted as a phase portrait in the v-u coordinate plane, which helps visualize the model’s behavior over time (Figure 8A and 8B). Non-linear neural models, such as the Izhikevich model, exhibit bifurcations (Figure 8C and 8D). These bifurcation points are crucial for describing the activity of the neural model. The excitatory neural units had a stable point when there was no external input (Figure 8A black circle). A stable point indicates that the neuron remains in a steady state, not generating action potentials. If the external input *I* exceeds the bifurcation point *I*=24.5 (Figure 8C) with the DG place input, the stable point is lost and bursting occurs (saddle-node bifurcation). The stable point reappeared when the external input decreased, and the firing became silent. When the inhibitory input was provided from the CA3 inhibitory unit, the quadric v-nullcline moved in the minus direction on the u-axis, causing a delay in firing (if the stable point did not appear) or stopping the firing altogether (if the stable point had appeared). These dynamics were the behavior of excitatory units in the simulated scenarios described above.

**Figure 8.**
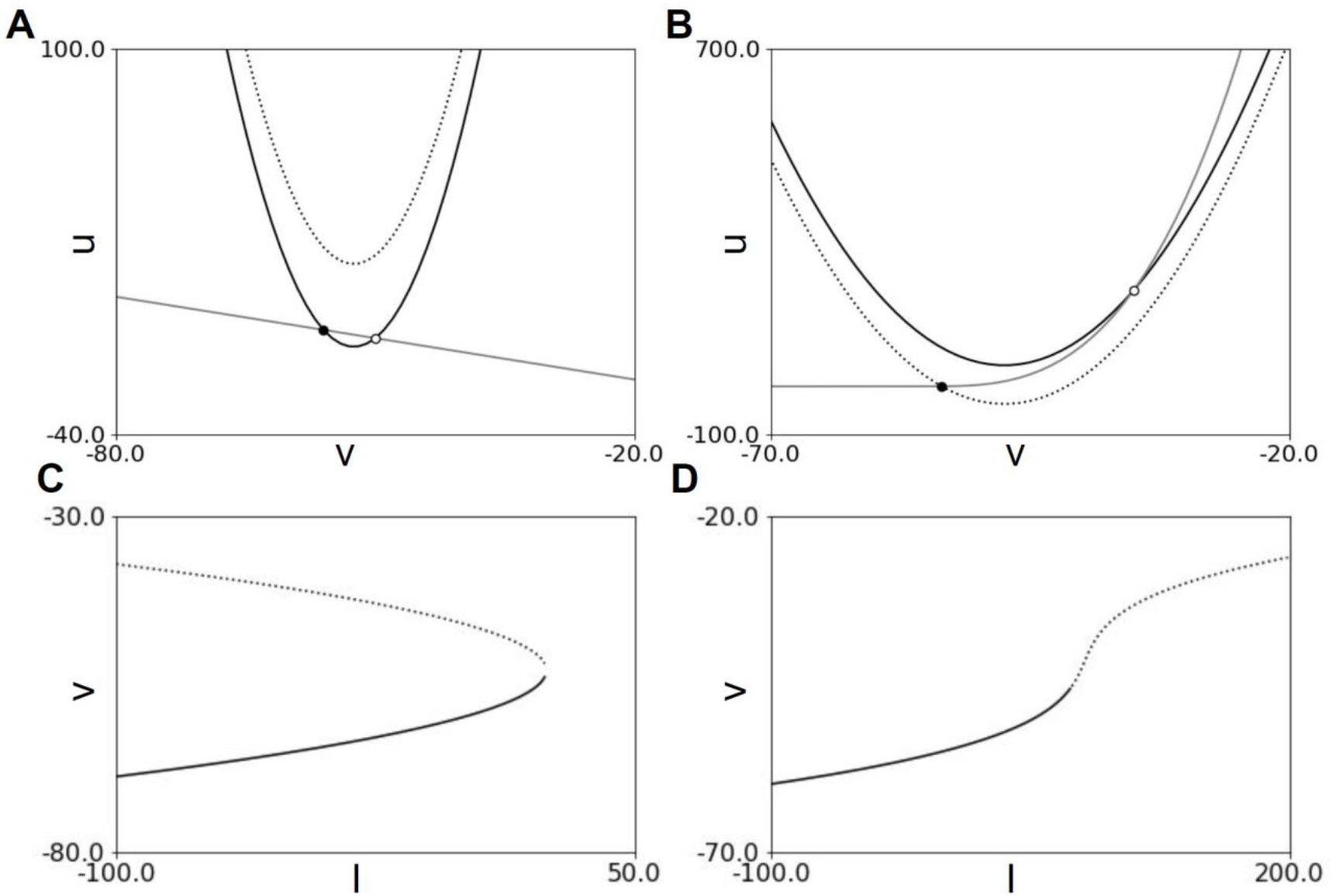
Bifurcation properties of excitatory and inhibitory units (A) Phase portrait of the excitatory unit. The black solid line indicates a v-nullcline without external input. The dotted line indicates v-nullcline with maximum I_p_. The gray line is the u-nullcline. The black and white circles indicate stable and unstable equilibrium, respectively. (B) Phase portrait of the inhibitory unit. The solid lines and black and white circles indicate the same as (A). The dotted line indicates v-nullcline with minimum I_th_. (C) Bifurcation diagram depending on external input *I* of the excitatory unit. The solid and dotted lines indicate stable and unstable equilibrium, respectively. (D) Bifurcation diagram depending on external input *I* of the inhibitory unit. The solid and dotted lines indicate the same as (C).

Without the MS theta inhibitory input (baseline, *I*=100), the inhibitory neural unit had an unstable point and exhibited bursting firing (Figure 8B solid black line and white circle). When the inhibitory theta input is received from the MS and the external input falls below *I*=73.7, the unstable point becomes stable (Andronov-Hopf bifurcation, Figure 8B dash black line, and black circle). When the point was stable, the dynamics of the unit converged on it and stopped firing. However, decreasing the amplitude of the theta inhibitory input caused the stable point to become unstable again, resulting in the reemergence of burst firing.

The amplitude of the external input affected the spike timing of the excitatory units in the theta cycle (Figure 9A–D). The MS inhibitory input controlled the theta cycle (Figure 9A). As described above, the CA3 excitatory unit does not spike without the external DG place input (Figure 8A solid line and Figure 9C c). When the external input amplitude increased slightly above the bifurcation point, the units began firing later in the theta cycle (Figure 9C a). With a small amplitude of the external input, the transition speed around the closest area between v- and u-nullclines is very slow (Figure 9C a lower). As the external input amplitude increased, the equilibrium disappeared, transition speed around the area increased, firing timing in the theta cycle started earlier, and the unit began to burst (Figure 9C b). The firings were restricted and reset by inhibitory inputs from the CA3 inhibitory unit, which spiked at specific phases in the theta cycle (Figure 9D). This resetting by the inhibitory input caused the start timings of the state in the v-u coordinate of each excitatory unit to be aligned with the theta cycle.

**Figure 9.**
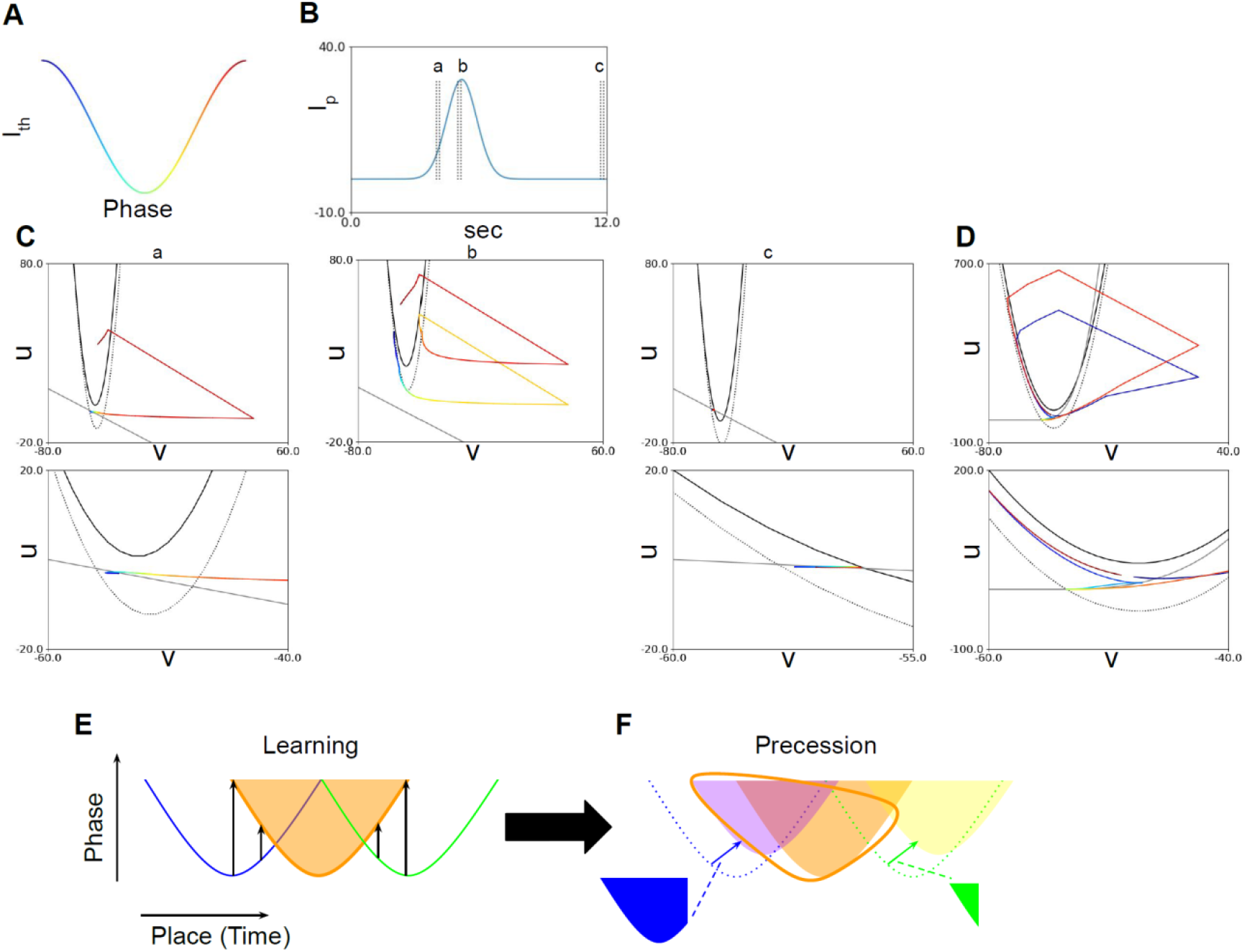
Firing timing in the theta cycle phase with place inputs (A) Correspondence between one cycle of theta oscillation and color in I_th_. (B) Single place input I_p_. Characters a, b, and c correspond to the one theta cycles plotted in (C). (C) The upper panels show the trajectory of the excitatory unit in the v-u phase portrait at the theta cycle of a, b, and c, respectively, in (B). Colors correspond to the colors of the theta cycle in (A). Lower panels are expansions of the upper panels around the closest or cross area between the v- and u-nullcline. (D) The upper panel shows the trajectory of the inhibitory unit without place input I_p_, corresponding to c in (B). The lower panel expands the upper panel around the closest area between the v- and u-nullcline. (E and F) Schematic illustration of this model’s acquisition process of phase precession and replay. (E) Before learning, the firing timings of a neuronal unit (orange shadow and line) on the phase-place (time) plane depend on place input strength, displaying a symmetric shape. Synaptic potentiation occurs (black arrow) with the CA3 excitatory unit responding to the anterior (blue line) and posterior (green line) place inputs. (F) The synaptic potentiation takes effect slightly later (colored arrows) than the firing of the previous neural unit (purple and yellow shadows), either accelerating the spike timing or inducing spikes in an additional theta cycle. The difference in plasticity-related factor accumulation (blue and green opaque shadows) and the overlapping area between the anterior (purple shadow) or posterior (yellow shadow) units and the center unit (orange shadow) results in a stronger effect from the anterior responding unit, leading to an asymmetric precession shape (orange line). Synaptic potentiation also affects replay.

As per the explanation, the theta-paced inhibitory input controls the spike timings of the CA3 excitatory units, but the phase precession pattern should be symmetric (no bias in the phase position of spikes before and after the peak position) as it depends solely on place input strength, which is symmetric in this model. Additional synaptic interactions shape the negative slope precession pattern. Before learning, the phase precession pattern of a CA3 excitatory neuronal unit is symmetric (Figures 3C and 9E orange shadow; referred to as the center unit). The learning process enhances recurrent synaptic weights, connecting the units responding to anterior and posterior place inputs (Figure 9E blue and green lines; referred to as anterior and posterior units, respectively) (Figure 2A). The difference in firing timing with the anterior and posterior units is also symmetric (Figure 9E arrows). The synaptic potentiation increases input amplitude and advances spike timing in the postsynaptic units (center unit) while also causing spikes in the late phase of the earlier theta cycle. The effect of synaptic input occurs slightly later than the presynaptic (anterior and posterior units) spike timings, creating an asymmetric effect where the anterior unit has more time to overlap with the activated timing of the center unit than the posterior unit (Figure 9F, purple-orange and yellow-orange shadow overlapping areas, respectively).

Furthermore, the anterior unit interacts with the center unit during the latter part of its activity, while the posterior unit interacts during the earlier part. This causes the plasticity-related factor (see Materials and methods) to accumulate more in the anterior unit, resulting in greater synaptic potentiation in the anterior unit than in the posterior unit (Figures 2C and 9F blue and green opaque shadows, respectively). The reason for the abnormal precession in unit #1 in Figure 3 is likely because there is no anterior unit for the unit #1. These mechanisms lead to an asymmetric phase precession pattern.

Additionally, the strengthened synapses induced sequential spikes by random noise input in the resting session. This mechanism established the theta phase precession in our models and replays after place learning.

## Discussion

In this study, we developed a neural circuit model based on anatomical evidence of the biological CA3 circuits. The model could learn and compress the order of external inputs from sparse DG projections using the Hebb rule and displayed theta phase precession and replay-like activity. The model could also perform additional learning and relearning. In this computational model, both phase precession and replay are generated by a shared underlying mechanism, namely synaptic enhancement of the CA3 recurrent connections. The synaptic modulation is governed by the interplay between theta rhythm inhibition and strength and temporal order of place inputs. These findings underscore the pivotal role of CA3 circuits in the acquisition of sequence information, significantly enriching our understanding of the mechanistic underpinnings of memory encoding processes.

The region responsible for the encoding sequence is still debated (Drieu and Zugaro, 2019). In this research, we assumed that the CA3 recurrent connection plays a role in sequence learning based on anatomical (the CA3 has a relatively dense recurrent connection) and electrophysiological (source of SWR) evidence (Buzsáki, 2015; Hájos et al., 2013; Rebola et al., 2017; Schlingloff et al., 2014). In most cases, replay in the SWR has been reported in the CA1 (Johnson and Redish, 2007; Karlsson and Frank, 2009). CA3 is likely the source of replay, as input from the CA3 has been observed when the CA1 ripple occurs (Buzsáki, 2015; Csicsvari et al., 2000) and the replay collapses when the CA3 is lesioned (Middleton and McHugh, 2016). The CA3 lesion also affects CA1 replay (Davoudi and Foster, 2019; Guan et al., 2021; Yamamoto and Tonegawa, 2017). There are also theta phase precession reports (Dragoi and Buzsáki, 2006; Fernández-Ruiz et al., 2017; Mizuseki et al., 2012; Oliva et al., 2016; Sharif et al., 2021) and reports of replay (Carr et al., 2012; Diba and Buzsáki, 2007; Hwaun and Colgin, 2019; Karlsson and Frank, 2009; O’Neill et al., 2008) in the CA3. The CA3-CA1 connection can also learn sequences, such as reservoir networks (Cazin et al., 2019; Enel et al., 2016). The DG-CA3 giant synapses are still a potential candidate for the region responsible for sequence learning. In this study, we excluded inputs from the entorhinal cortex, which have been shown to affect theta phase precession (Fernández-Ruiz et al., 2017; Schlesiger et al., 2015) and replay (Chenani et al., 2019; Yamamoto and Tonegawa, 2017) in the hippocampus. To fully understand the main region responsible for sequence learning, integrating these experimental findings and computational frameworks will be necessary in the future.

Models that explain theta phase precession can be divided into three main categories (Drieu and Zugaro, 2019): detuned oscillators model (Geisler et al., 2010; O’Keefe and Recce, 1993), somato-dendritic interference model (Harris et al., 2002; Kamondi et al., 1998; Mehta et al., 2002), and network connectivity model (Jensen and Lisman, 1996; Romani and Tsodyks, 2015; Skaggs et al., 1996; Tsodyks et al., 1996). The detuned oscillators model posits that phase precession arises from single place cell oscillating at a slightly faster frequency than the population signal (LFP) theta oscillation, with both oscillators gradually shifting due to their frequency difference. Our findings do not support this model, as the LFP (summation of all excitatory membrane potential) and individual neuronal firing or membrane potential frequency in our model were identical (Figure 3B).

In contrast, the somato-dendritic interference model suggests that a combination of oscillatory somatic inhibition and transient ramp-like dendritic excitation determines action potential timing. Our results support this model, as place input strength dictates spike timing in space without recurrent inputs (Figures 8 and 9). Furthermore, the expansion of the place fields (Figure 1D–F) also supports developing the precession with the somato-dendritic interference model (Mehta et al., 2002, 2000). Network connectivity models propose that phase precession results from transmission delays between asymmetrically connected cells (Jensen and Lisman, 1996; Romani and Tsodyks, 2015; Skaggs et al., 1996; Tsodyks et al., 1996). Our results align with those obtained in these models, as our model exhibited additional spikes in the late phase of the early theta cycle after learning (Figures 1–3). These findings highlight the potential roles of somato-dendritic interference and network connectivity in phase precession, although our model does not exhaustively explore these mechanisms.

In our model, phase precession gradually forms through learning, consistent with the results of Mehta et al. (2002). However, this does not align with the results of Feng et al. (2015), who reported that phase precession of individual place cells was present from the initial trial and became more aligned across cells as the number of trials increased, forming theta sequences. In this study, our model assumed nearly uniform membrane properties and synaptic connections in its initial state. By introducing variabilities in these values, such as distinct cellular properties and prewired connections for each cell, it might be possible to reproduce the results of Feng et al. (2015) within our model. Another possibility is that the difference arises from the distinct characteristics of CA3 and CA1, as these reports focus on CA1.

There is a debate regarding the replay, i.e., whether it is pre-existing (Dragoi and Tonegawa, 2011) or acquired (Silva et al., 2015). Many computational mechanisms for achieving replay sequences have been proposed in computational modeling studies (August and Levy, 1999; Ecker et al., 2022; Haga and Fukai, 2018; Jahnke et al., 2015; Jensen et al., 1996; Nicola and Clopath, 2019; Reifenstein et al., 2021). The results of this study explained how a recurrent network could acquire the replay sequences, incorporating both pre-existing and acquired aspects. Specifically, task-related and strongly connected synapses before learning tend to be enhanced through repeated experience.

In our model, primarily forward replay was observed. However, it may be possible to reveal different patterns, such as reverse replay, by altering synaptic learning rules (for example, our model used an asymmetric Hebbian rule, but other options could include standard STDP [Bi and Poo, 1998] or symmetric STDP [Mishra et al., 2016]) or by imposing of an upper limit on synaptic strength.

Models similar to our model include those designed by Jensen et al. (1996) (Jensen et al., 1996; Jensen and Lisman, 1996), and Ecker et al. (2022). Both ours and these other models learned the order in which place cells are active in the CA3 recurrent connection through the Hebb rule or similar learning rules (symmetric STDP) (Mishra et al., 2016). As a result of learning, all models showed a non-Gaussian, asymmetric distribution of synapse weights. Experimental results in biological brains also suggest the occurrence of similar phenomena (Choi et al., 2018). However, several distinctive features of our model set it apart: (1) In contrast to the other models, our model included theta input as inhibitory input to inhibitory units. This is supported by optogenetic experiments showing that theta rhythm input is mediated by inhibitory projections from MS to PV-positive inhibitory neurons in the hippocampus (Zutshi et al., 2018). (2) Our model also reproduced the decrease in inhibitory neuron activity during sleep or rest (Alfonsa et al., 2022; Miyawaki and Diba, 2016; Mizuseki and Buzsáki, 2013), which leads to increased random firing and the potential induction of replay during resting sessions. (3) Jensen et al. (1996) focused on phase precession (with no mention of replay), and Ecker et al. (2022) demonstrated the occurrence of SWR-like activity and the accompanying forward and reverse replay (with no mention of precession). Meanwhile, our model aims to reproduce both the characteristic precession and replay activity in the hippocampus through learning in the CA3 local circuit and recurrent connection. These unique aspects of our model provide new insights and a more comprehensive understanding of the CA3 circuit’s role in theta phase precession and replay activity in the hippocampus.

The primary limitations of this study arise from addressing only the CA3 region’s minimal elements, such as the recurrent connectivity and plasticity in CA3, place input from the DG, and inhibitory theta rhythms from the MS. This focus resulted in a less robust depiction of several phenomena, such as phase precession in both slope and range, and a dominance of forward-ordered replay, compared to empirical biological observations. The alignment with experimental data could potentially be improved by incorporating elements such as inputs from the EC (Fernández-Ruiz et al., 2017; Hafting et al., 2008; Schlesiger et al., 2015) or CA3 inhibitory recurrent connections (Ecker et al., 2022; Tiesinga and Sejnowski, 2009), alternative synaptic plasticity rules, and variability in neuronal or synaptic properties.

In this study, we proposed a shared mechanism, synaptic enhancement, for compressing the sequence of external events into theta phase precession and replay. The synaptic modulation is governed by the interaction between theta inhibitory and place excitatory inputs. However, several points still need to be elucidated. Is the formation of precession and replay pre-existing or acquired? How does the neuronal synchronization state of the hippocampus change during activity and rest? How does the replay occur? Does replay in CA3 have different functions than replay in the CA1 and other brain regions? Is synaptic plasticity activated during replay? What are the contributions of circuits other than CA3? What are the effects of learning on replay? Are the replayed sequences varied and evaluated during each replay for abstraction, planning, and inference? How do these sequences interact with cortical networks to facilitate their functions? Further research is needed to investigate these questions from experimental and computational approaches.

## Materials and methods

### Small model

A small model is utilized to deeply analyze the dynamics of the system under well-controlled conditions. The CA3 model consisted of eight excitatory and one inhibitory unit (Figure 1A). The excitatory units received the DG place inputs, recurrent CA3 connections, and inhibitory inputs from the CA3 inhibitory unit. The inhibitory unit received the theta MS inhibitory input and excitatory inputs from the CA3 excitatory units. The activity of the units was modeled using the Izhikevich model (Izhikevich, 2010), and the parameters were chosen according to sections “8.4.1 Hippocampal CA1 Pyramidal Neurons” and “8.2.6 Fast Spiking Interneurons” for the excitatory and inhibitory units, respectively.

The Izhikevich model is described as follows:

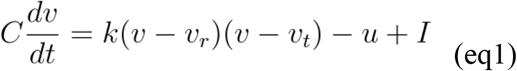

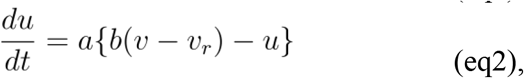

where *v* is the fast membrane potential, *u* is the slow recovery variable that allows for the replication of various neural cell types, *v*_*r*_ is the resting potential, *v*_*t*_ is the threshold potential for spike initiation, *C* is the membrane capacitance, *a* is the recovery time constant of *u, k* and *b* are scaling coefficients, *I* is the input to the neural unit, and *v*_*peak*_ is the peak potential; if *v* exceeds *v*_*peak*_, *v* and *u* are reset to *v*→ *c* and *u* → *u* + *d*, respectively. In the inhibitory unit,

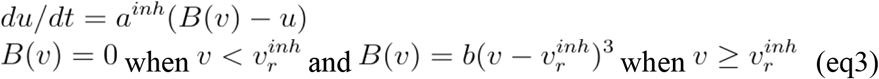

(Chap. 8.2.6, Izhikevich, 2010).

The CA3 excitatory unit receives excitatory DG place inputs, recurrent CA3 connections, and inhibitory inputs from the CA3 inhibitory unit.

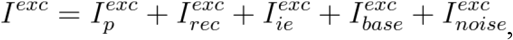

where 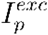 is the place input from the DG, 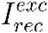 is the recurrent input from the CA3 excitatory units, 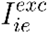 is the input from CA3 inhibitory unit to CA3 excitatory unit, 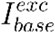 is a constant baseline input, and 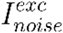 is a Gaussian noise input with 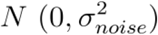. Similarly, the CA3 inhibitory unit receives excitatory inputs from CA3 excitatory units and inhibitory theta input from the MS.

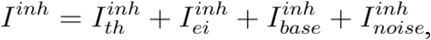

where 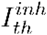 is the theta input from MS, 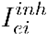 is input from the CA3 excitatory units, 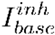 is a constant baseline input, and 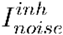 is a Gaussian noise input. In the simulation, the *<*3;_*noise*_ is scaled by the square of the time step 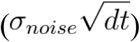.

The synaptic inputs 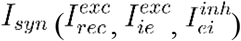 of the unit were described as follows:

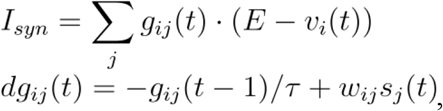

where *g*_*ij*_ (t) is a synaptic conductance between presynapse *j* and postsynapse *i, E* is an equilibrium potential (*E*_*ampa*_ = 0 for excitatory synapses and *E*_*gapa*_ = −65 for inhibitory synapses), *τ* is a time decay constant of the synaptic conductance *(τ*_*ampa*_ = 10 ms, *τ*_*gapa*_ = 20 ms), *s*_*j*_ (t) is a spike (0 or 1) of the presynaptic neural unit, and *w*_*ij*_ is a synaptic weight between pre and postsynaptic neural units. Notably, 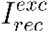 and 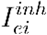 represent excitatory synapses (AMPA) and 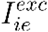 represents inhibitory synapses (GABA-A).

The DG-CA3 excitatory unit connection (place input) was assumed to be sparse, meaning that each CA3 excitatory unit received only one place input. The amplitude of the place inputs was set high enough to induce spikes in the CA3 excitatory units with only one place input. This model used four place inputs for each session in eight DG inputs. The CA3-CA3 recurrent connections were all-to-all and plastic connections (please refer to Plasticity section). All CA3 excitatory units sent spikes to the CA3 inhibitory unit, which in turn sent feedback inhibition to all CA3 excitatory units.

### External inputs

In this model, we assumed that the place inputs were from DG to CA3 excitatory neural units, and the theta inputs were from MS to CA3 inhibitory units.

The place inputs from DG were modeled as follows:

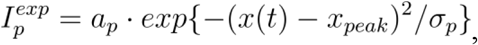

where *a*_*p*_ is the amplitude of the place input, *x* (*t*) is the current position of the agent, *x*_*peak*_ is the peak position of the place field, and the *σ*_*p*_ is the width of the place field. The agent’s speed is assumed to be constant in this simulation; thus, *x* (*t*) = *t* cm (velocity is 1 cm/s). The input orders were as follows: initial learning, units #1-2-3-4; additional learning, units #4-5-6-7; and relearning, units #8-1-3-6.

The theta oscillation inputs from MS were modeled as follows:

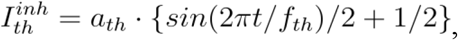

where *a*_*th*_ is the amplitude of the theta input and *f*_*th*_ is the frequency of the theta oscillation. As the agent’s speed is assumed to be constant in this simulation, *f*_*th*_ was also constant.

### Plasticity

The synaptic weights of the CA3-CA3 recurrent connection were plastically changed according to Hebbian and regularization rules (Kim et al., 2020; Weber and Sprekeler, 2018). Plasticity-related factors, such as calcium ion and calmodulin-dependent kinase II, are assumed to have a longer time constant than synaptic inputs (Rogerson et al., 2014). Activity of the plasticity-related factor was modeled as pulse activation and exponential decay, the same as synaptic activities (Jensen et al., 1996). Furthermore, we implemented a superlinear coincident detection; when two inputs from different sources arrive simultaneously, the response is much higher than the simple summation of the responses to these two inputs (Brandalise and Gerber, 2014; London and Häusser, 2005).

The Hebbian plasticity between the postsynapse unit *i* and the presynapse unit *j* was modeled as:

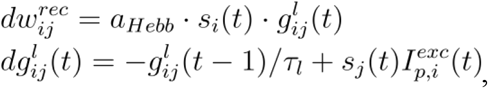

where *a*_*Hepp*_ is a learning rate of the Hebbian learning, *s*_*i*_ and *s*_*j*_ are spikes of postsynaptic and presynaptic neural units, respectively, and 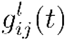 is the assumed activity of plasticity-related factors, which has similar dynamics to the synaptic conductance *g*_*ij*_ (*t*) but with a longer decay time constant *τl* than that with voltage change. The coincident detection was implemented by multiplying the place cell input 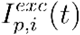 from DG with *s*_*j*_ (*t*). Self-connections were excluded, meaning that 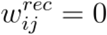 (if *i* = *j*).

The regularization rule was modeled as:

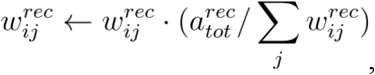

where 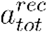is the gain of the regularization, by which the regularization rule ensures that the summation of the postsynaptic weights is permanently restricted.

The parameter values used in this model are summarized in Table 1.

**Table 1.** Parameters of the small CA3 model (values are for Figures 1–4)

| Parameter name | Symbol | Value | Parameter name | Symbol | Value |
| --- | --- | --- | --- | --- | --- |
| Number of trials | $N_{trial}$ | 10 | <u>Synapse</u> | | |
| Time of one trial | $T$ | 12000 ms | Reversal potential (Exc) | $E_{ampa}$ | 0 |
| Time derivative | $dt$ | 0.5 ms | Reversal potential (Inh) | $E_{gaba}$ | -65 |
| <u>CA3 excitatory neural unit</u> | | | Time constant (Exc) | $\tau_{ampa}$ | 10 ms |
| Number of units | $N^{exc}$ | 8 | Time constant (Inh) | $\tau_{gaba}$ | 20 ms |
| Parameters of the Izhikevich Model | $C^{exc}$ | 50 | Synaptic weight (CA3 Inh → CA3 Exc) | $w^{ie}$ | 1 |
| | $k^{exc}$ | 0.5 | Synaptic weight (CA3 Exc → CA3 Inh) | $w^{ei}$ | 1 |
| | $v_r^{exc}$ | -60 | Synaptic weight (DG j → CA3 Exc i) | $w^{jc}$ | 1 ($i = j$) or 0 ($i \neq j$) |
| | $v_t^{exc}$ | -45 | Synaptic weight (MS → CA3 Inh) | $w^{ti}$ | 1 |
| | $v_{peak}^{exc}$ | 40 | <u>External Input</u> | | |
| | $a^{exc}$ | 0.02 | Amplitude of place input | $a_p$ | 30 |
| | $b^{exc}$ | -0.5 | Peak position of place input | $x_{peak}$ | 2, 3, 4, 5 cm |
| | $c^{exc}$ | -45 | Window width of place input | $\sigma_p$ | 1 cm |
| | $d^{exc}$ | 50 | Amplitude of theta input | $a_{th}$ | -80 |
| <u>CA3 Inhibitory neural unit</u> | | | Frequency of theta input | $f_{th}$ | 7 Hz |
| Number of units | $N^{inh}$ | 1 | Baseline of excitatory input | $I_{base}^{exc}$ | 20 |
| Parameters of the Izhikevich Model | $C^{inh}$ | 20 | Baseline of inhibitory input | $I_{base}^{inh}$ | 100 |
| | $k^{inh}$ | 1 | Noise level of excitatory input | $\sigma_{noise}^{exc}$ | 0 |
| | $v_r^{inh}$ | -55 | Noise level of inhibitory input | $\sigma_{noise}^{inh}$ | 0 |
| | $v_t^{inh}$ | -40 | <u>Plasticity</u> | | |
| | $v_{peak}^{inh}$ | 25 | The learning rate for Hebbian plasticity | $a_{Hebb}$ | 0.0083 (100/T) |
| | $a^{inh}$ | 0.2 | The synaptic time constant for plasticity-related factor | $\tau_l$ | 300 ms |
| | $b^{inh}$ | 0.025 | Summation limit for regularization | $a_{tot}^{rec}$ | 1 |
| | $c^{inh}$ | -45 | Velocity in place | — | 1 cm/s |
| | $d^{inh}$ | 20 | | | |

CA, cornu ammonis; DG, dentate gyrus; Exc, Excitatory; Inh, Inhibitory; MS, medial septum.

### Resting session

In the resting session, it is feasible to assume the absence of place input, given that the animal is not in the place of running session, as well as the absence of theta input, given that the animal is not running, concurrently with the presence of spontaneous activity, including replay (Buzsáki, 2015; Drieu and Zugaro, 2019). Neural inhibition is weakened during sleep or rest (Alfonsa et al., 2022; Miyawaki and Diba, 2016; Mizuseki and Buzsáki, 2013). Thus, we removed the external place and theta inputs and added independent noise inputs into each CA3 excitatory unit to induce spontaneous spiking. In addition, the synaptic plasticity was also turned off. The specific parameter changes for the resting session are as follows:

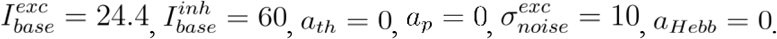

### Altered models

For the model with self-connections in the recurrent synapse (Figure 4—figure supplement 1), we set 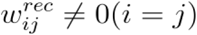, which were excluded in the original model.

For the model without coincident detection (Figure 4—figure supplement 2), we replaced 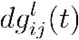 with

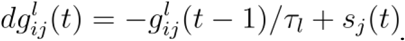

We set *a*_*Hepp*_ =0.083 (10 times larger) to gain a higher learning rate (Figure 4—figure supplement 2 F–J).

For the model with a short learning time constant (Figure 4—figure supplement 3), we set *τ*_*l*_ = 20 ms and set *a*_*Hepp*_ = 0.083 to gain a higher learning rate (Figure 4—figure supplement 3 F–J).

We set *a*_*th*_ = 0 and 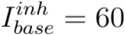for the model without theta input from the MS (Figure 4—figure supplement 4).

Other settings and processes were the same as in the small CA3 model (Figures 1–4).

### Large model

To analyze dynamics under conditions that are more similar to the actual brain, specifically with sparce stochastic connections, we utilized a larger model. In the large model, the synaptic connections between the CA3 circuit units were modeled as probabilistic, indicating that the connections in the CA3 circuit are not predetermined (Rebola et al., 2017). The large model was simulated using the Brian 2 neural simulator (Stimberg et al., 2019). The number of CA3 excitatory units was *N*_*exc*_ = 800, and the number of CA3 inhibitory units was *N*_*inh*_ = 200. The probability of the DG (place input)-CA3 excitatory connection was (Claiborne et al., 1986; Rebola et al., 2017), CA3 excitatory-CA3 recurrent excitatory connection was *p*_*pe*_ = 0.005 (Guzman et al., 2016), CA3 inhibitory-CA3 excitatory connection was *p*_*ie*_ = 0.25, CA3 excitatory-CA3 inhibitory connection was *p*_*ei*_ =0.25, and MS (theta input)-CA3 inhibitory connection was *p*_*ti*_ = 1. These probabilities were based on the results of Ecker et al. (2022) and Rebola et al. (2017). The synaptic weights were adjusted with 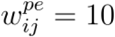 (if connected) and 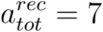. There were three trials in the running session, and the resting session was performed before and after the running session. Other parameters were the same as in the small model.

In Figure 7H and L, 15 models were executed with different random seeds. The units for the phase precession analysis were selected according to two criteria: they responded to the place input. They exhibited an expansion of firing through learning (more than 0.2 s earlier in the last trial than in the first).

### Theta phase precession

The LFP was calculated as the summation of the membrane potentials of all excitatory units. Next, the LFP signal was filtered using a bandpass filter (5–12 Hz), and a Hibert transform was applied. The phase component was extracted from the transformed signal. Finally, a linear regression analysis was used to plot the place vs. phase points to estimate the slope value.

### Crosscorrelogram

The spike timing differences were collected for each unit pair (unit *I* and unit *j, i* ≠ *j*). These timing differences were grouped according to the activation status of the units during the running session and whether both units were responded units, one unit in the pair was a responded unit while the other was not, or both units were non-responded units. In the analysis of additional learning and relearning, pairs of initial and new sequence sets were also plotted. The densities were calculated over a range of -600 to 600 ms.

### Replay sequence

Linear regression was performed between the difference in peak positions of place inputs during learning and lags during replay after learning, with a lag range from -200 to 200 ms of pairs between the responded units. The unit that responded to the first-place input was excluded.

### Dynamical analysis

The nullcline lines were defined as the points where the formulas (eq1) and (eq2) (for excitatory, eq3 instead of eq2 for inhibitory) were equal to 0. The equations represented the nullclines:

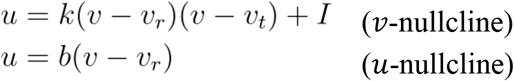

The equilibrium points were determined as the intersection between the -nullcline and the -nullcline. The stability of each equilibrium point was determined by the eigenvalues of the Jacobian matrix of the system represented by the formulas (eq1) and (eq2). If the two eigenvalues are both negative, the equilibrium point is stable. Otherwise, it is unstable (Izhikevich, 2010). When the formulas (eq1) and (eq2) were redefined as follows:

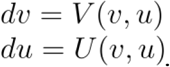

the Jacobian matrix,, was defined as:

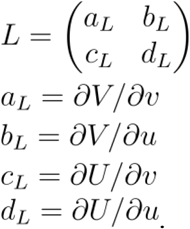

The eigenvalues *λ*, and eigenvectors *x*, of the Jacobian matrix *L*, can be computed by solving the equation:

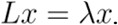

The eigenvalues can be obtained by solving the following equation:

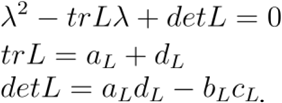

Both eigenvalues should be negative for the equilibrium point to be stable, corresponding to *trL <* 0 and *detL* > 0. If the conditions are not met, the equilibrium point is unstable.

The equilibrium points of excitatory units 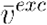 were obtained by solving the following equation:

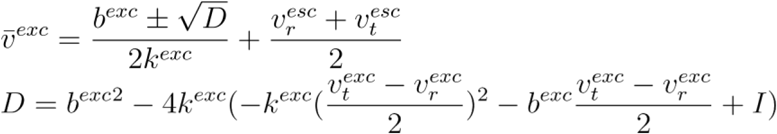

It exhibited a saddle-node bifurcation when the constant input current *I* was <24.5. In this case, the negative side equilibrium was stable, while the positive side was unstable (Izhikevich, 2010). The equilibrium points disappeared at above =24.5.

The equilibrium points of inhibitory units 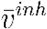 were obtained by solving for *v* in following equation:

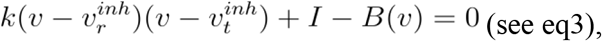

which was solved using the scipy.optimize.fsolve method with a moving constant *I*. The dynamics of the inhibitory unit exhibited the Andronov–Hopf bifurcation depending on *I*. Then,

*tr L*^*inh*^ and *det L*^*inh*^ were

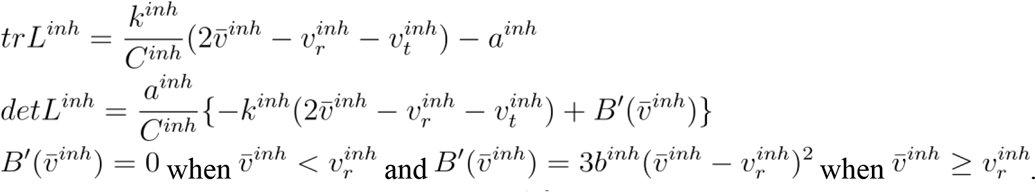

The equilibrium point was stable when *tr L*^*inh*^ ≤ 0 and unstable when *tr L*^*inh*^ > 0, as *det L*^*inh*^ is always positive. The bifurcation point occurred at *I* =73.7.

## Code Availability Statement

The scripts for this study can be found in the Google Colaboratory https://colab.research.google.com/drive/1mwl6ti3lAFtVwtfoZBPhdbSUXrUoCd7v?usp=sharing.

## Acknowledgments

We thank Hiroyuki Miyawaki and Takuya Isomura for their valuable contributions to the discussion. This work was supported by JSPS KAKENHI (23H02788 to K.M.).

---

**Figure 4 —figure supplement 1.**
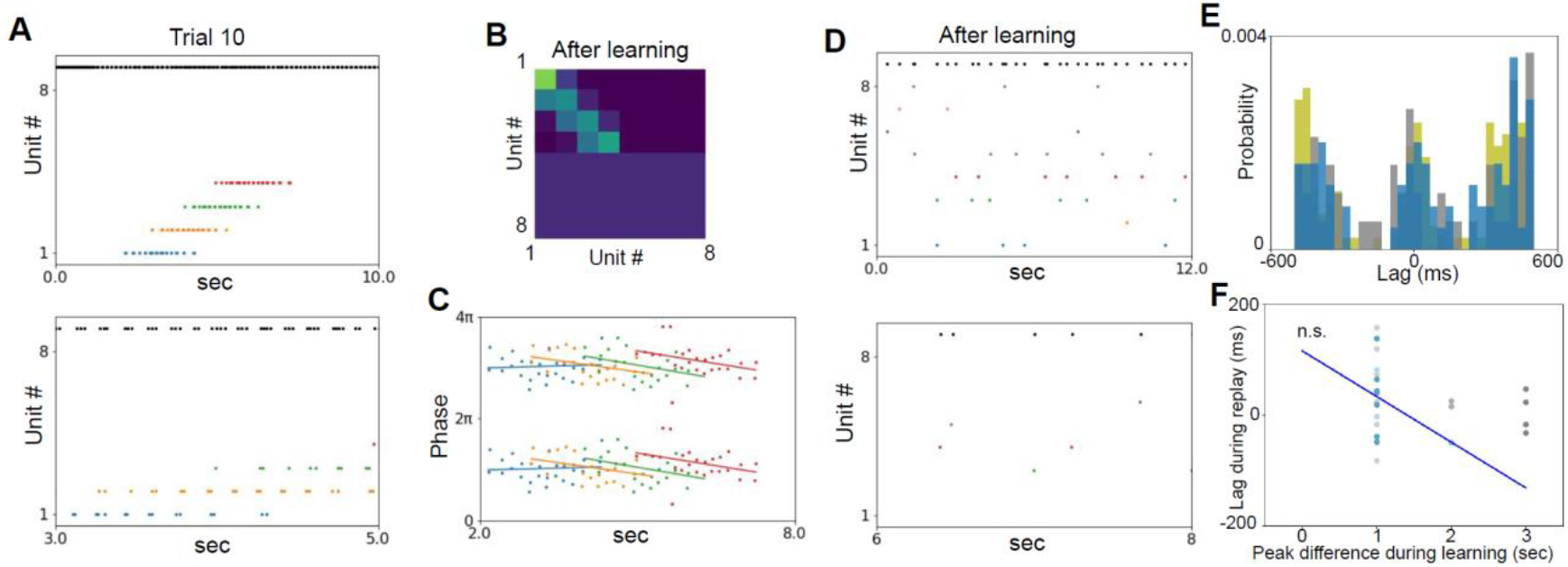
Small model with self-connection in the recurrent synapse It exhibits synaptic weight enhancement with neighbor units and theta phase precession but not the replay. (A) Upper: spike plot of trial 10. Bottom: zoom in for 3 to 5 s of the upper panel. (B) Synapse weights after learning. (C) Phase precession after learning. (D) Upper: spike activity in the resting session with noise input after learning. Bottom: 2 s zoom in of the upper panel. (E) Crosscorrelogram of the resting session after learning. (F) Scatter plot of the difference in peak positions of place inputs during learning and lags during replay after learning between responded units (linear regression: slope=-82, intercept=115, r=-0.42, p=0.22, n=10). The colors and initial setting correspond to those in Figures 1–4. n.s. p > 0.05

**Figure 4 —figure supplement 2.**
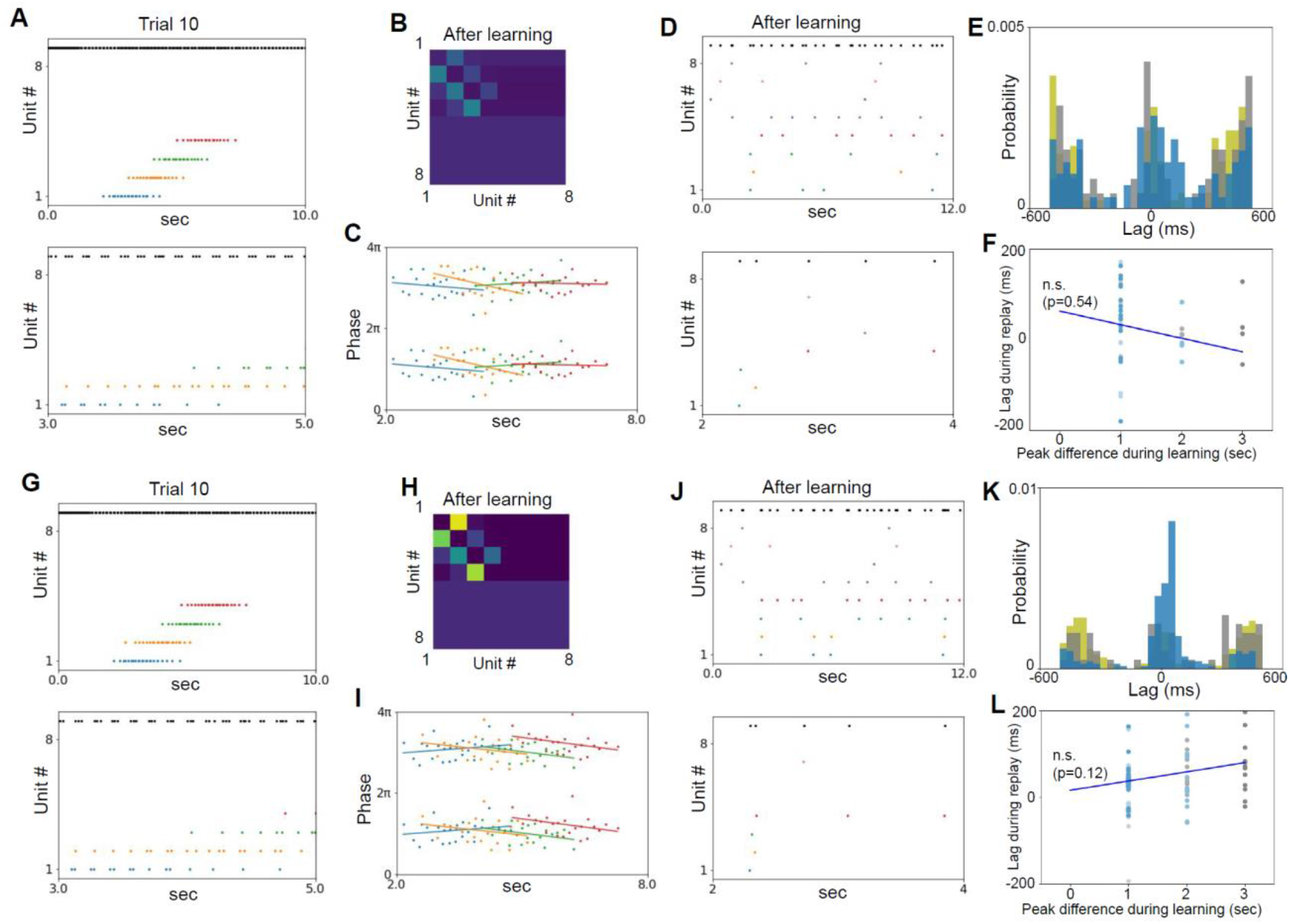
Small model without coincident detection The synaptic weight enhancement has weakened. The theta phase precession and the replays were not observed, which were recovered with a large learning rate. (A) Upper: spike plot of trial 10. Bottom: zoom in for 3 to 5 s of the upper panel. (B) Synapse weights after learning. (C) Phase precession after learning. (D) Upper: spiking activity in resting session with noise input after learning. Bottom: 2 s zoom in of the upper panel. (E) Crosscorrelogram of the resting session after learning. (F) Scatter plot of the difference in peak positions of place inputs during learning and lags during replay after learning between responded units (linear regression: slope=-30, intercept=4, r=-0.12, p=0.54, n=28). (G–L) Same as the above panels with a 10 times larger learning rate (L, linear regression: slope=21, intercept=16, r=0.19, p=0.12, n=72). The colors and initial setting correspond to those in Figures 1–4. n.s. p > 0.05

**Figure 4 —figure supplement 3.**
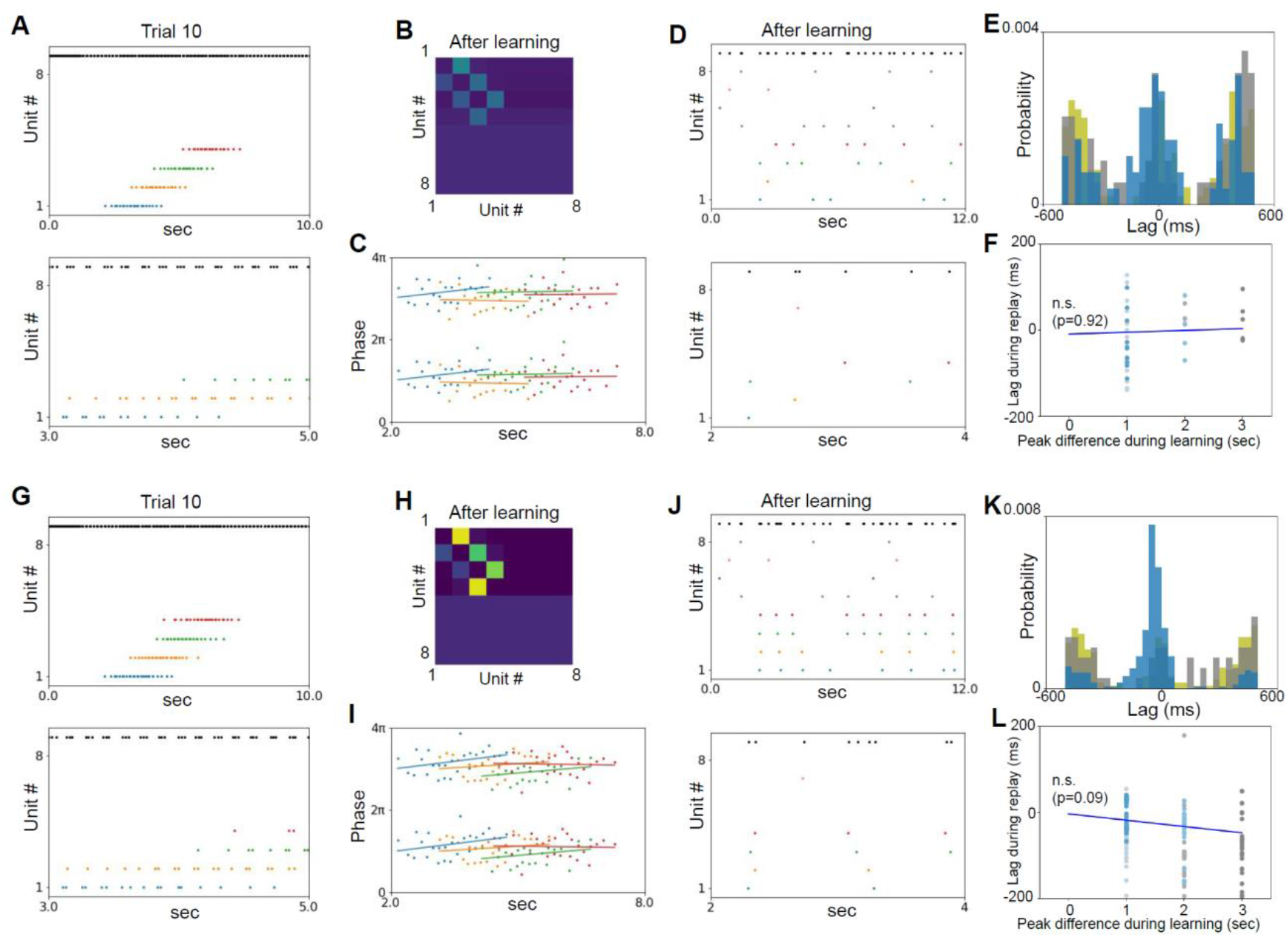
Small model with a short time constant of plasticity-related factor The synaptic weight enhancement has weakened. The theta phase precession and the replays were not observed, which were recovered with a larger learning rate while the order was reversed. (A) Upper: spike plot of trial 10. Bottom: zoom in for 3 to 5 s of the upper panel. (B) Synapse weights after learning. (C) Phase precession after learning. (D) Upper: spiking activity in resting session with noise input after learning. Bottom: 2 s zoom in of the upper panel. (E) Crosscorrelogram of the resting session after learning. (F) Scatter plot of the difference in peak positions of place inputs during learning and lags during replay after learning between responded units (linear regreassion: slope=4.2, intercept=-9.5, r=0.023, p=0.92, n=21). (G–L) Same as the above panels with a 10-fold higher learning rate (L, linear regression: slope=-15, intercept=-3.6, r=-0.15, p=0.094, n=128). The colors and initial setting correspond to those in Figures 1–4. n.s. p > 0.05

**Figure 4 —figure supplement 4.**
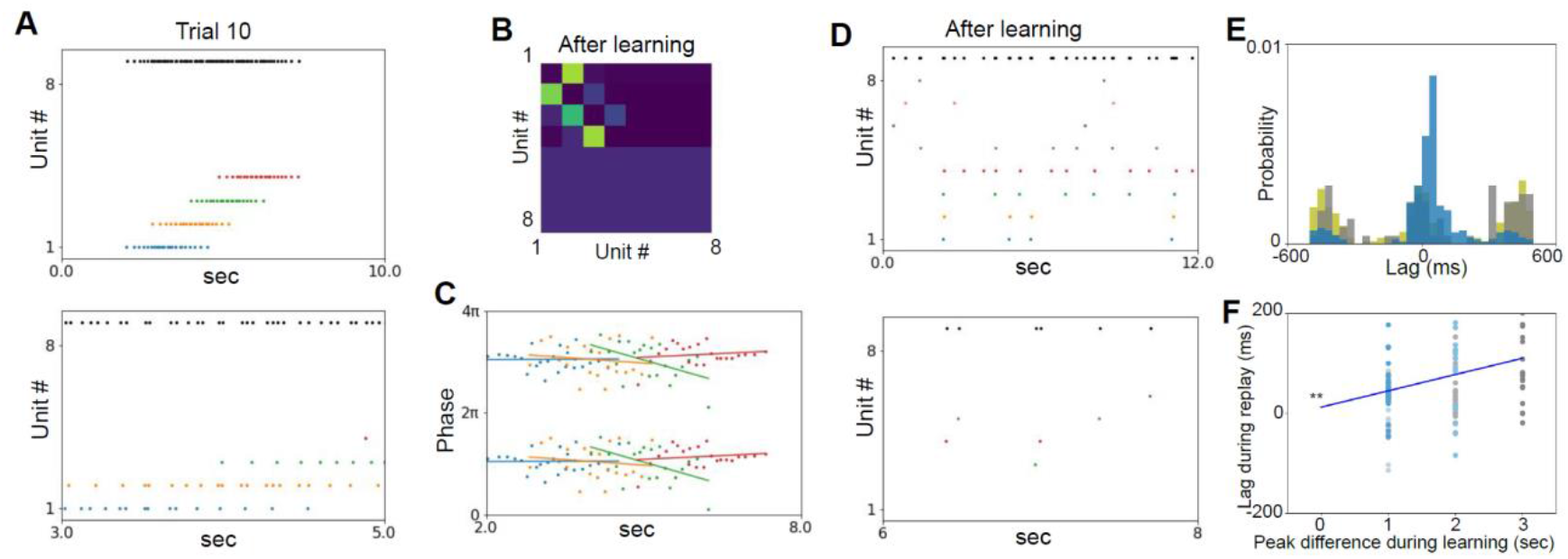
Small model without theta input from MS The theta phase precession is not observed, but the synaptic weight enhancement with neighboring units and the relay is still observed. (A) Upper: spike plot of trial 10. Bottom: zoom in for 3 to 5 s of the upper panel. (B) Synapse weights after learning. (C) Phase precession after learning. (D) Upper: spiking activity in resting session with noise input after learning. Bottom: 2 s zoom in of the upper panel. (E) Crosscorrelogram of the resting session after learning. (F) Scatter plot of the difference in peak positions of place inputs during learning and lags during replay after learning between responded units (linear regression: slope=33, intercept=12, r=0.26, p=0.009, n=100). The colors and initial setting correspond to those in Figures 1–4. MS, medial septum. ** p < 0.01

## References

Alfonsa H, Burman RJ, Brodersen PJN, Newey SE, Mahfooz K, Yamagata T, Panayi MC, Bannerman DM, Vyazovskiy VV, Akerman CJ. 2022. Intracellular chloride regulation mediates local sleep pressure in the cortex. Nat Neurosci. doi:10.1038/s41593-022-01214-2

Andersen P, Morris R, Amaral D, Bliss T, O’Keefe J, editors. 2006. The Hippocampus Book. Oxford University Press. doi:10.1093/acprof:oso/9780195100273.001.0001

August DA, Levy WB. 1999. Temporal sequence compression by an integrate-and-fire model of hippocampal area CA3. J Comput Neurosci 6:71–90. doi:10.1023/a:1008861001091

Banino A, Barry C, Uria B, Blundell C, Lillicrap T, Mirowski P, Pritzel A, Chadwick MJ, Degris T, Modayil J, Wayne G, Soyer H, Viola F, Zhang B, Goroshin R, Rabinowitz N, Pascanu R, Beattie C, Petersen S, Sadik A, Gaffney S, King H, Kavukcuoglu K, Hassabis D, Hadsell R, Kumaran D. 2018. Vector-based navigation using grid-like representations in artificial agents. Nature 557:429–433. doi:10.1038/s41586-018-0102-6

Bi GQ, Poo MM. 1998. Synaptic modifications in cultured hippocampal neurons: dependence on spike timing, synaptic strength, and postsynaptic cell type. J Neurosci 18:10464–10472. doi:10.1523/JNEUROSCI.18-24-10464.1998

Brandalise F, Gerber U. 2014. Mossy fiber-evoked subthreshold responses induce timing-dependent plasticity at hippocampal CA3 recurrent synapses. Proc Natl Acad Sci U S A 111:4303–4308. doi:10.1073/pnas.1317667111

Buzsáki G. 2015. Hippocampal sharp wave-ripple: A cognitive biomarker for episodic memory and planning. Hippocampus 25:1073–1188. doi:10.1002/hipo.22488

Buzsáki G. 1986. Hippocampal sharp waves: their origin and significance. Brain Res 398:242–252. doi:10.1016/0006-8993(86)91483-6

Buzsáki G, Leung LW, Vanderwolf CH. 1983. Cellular bases of hippocampal EEG in the behaving rat. Brain Res 287:139–171. doi:10.1016/0165-0173(83)90037-1

Buzsáki G, Moser EI. 2013. Memory, navigation and theta rhythm in the hippocampal-entorhinal system. Nat Neurosci 16:130–138. doi:10.1038/nn.3304

Carr MF, Karlsson MP, Frank LM. 2012. Transient slow gamma synchrony underlies hippocampal memory replay. Neuron 75:700–713. doi:10.1016/j.neuron.2012.06.014

Cazin N, Llofriu Alonso M, Scleidorovich Chiodi P, Pelc T, Harland B, Weitzenfeld A, Fellous J-M, Dominey PF. 2019. Reservoir computing model of prefrontal cortex creates novel combinations of previous navigation sequences from hippocampal place-cell replay with spatial reward propagation. PLoS Comput Biol 15:e1006624. doi:10.1371/journal.pcbi.1006624

Chadwick A, van Rossum MC, Nolan MF. 2016. Flexible theta sequence compression mediated via phase precessing interneurons. Elife5. doi:10.7554/eLife.20349

Chenani A, Sabariego M, Schlesiger MI, Leutgeb JK, Leutgeb S, Leibold C. 2019. Hippocampal CA1 replay becomes less prominent but more rigid without inputs from medial entorhinal cortex. Nat Commun 10:1341. doi:10.1038/s41467-019-09280-0

Choi J-H, Sim S-E, Kim J-I, Choi DI, Oh J, Ye S, Lee J, Kim T, Ko H-G, Lim C-S, Kaang B-K. 2018. Interregional synaptic maps among engram cells underlie memory formation. Science 360:430–435. doi:10.1126/science.aas9204

Claiborne BJ, Amaral DG, Cowan WM. 1986. A light and electron microscopic analysis of the mossy fibers of the rat dentate gyrus. J Comp Neurol 246:435–458. doi:10.1002/cne.902460403

Csicsvari J, Hirase H, Mamiya A, Buzsáki G. 2000. Ensemble patterns of hippocampal CA3-CA1 neurons during sharp wave-associated population events. Neuron 28:585–594. doi:10.1016/s0896-6273(00)00135-5

Davoudi H, Foster DJ. 2019. Acute silencing of hippocampal CA3 reveals a dominant role in place field responses. Nat Neurosci 22:337–342. doi:10.1038/s41593-018-0321-z

Diba K, Buzsáki G. 2007. Forward and reverse hippocampal place-cell sequences during ripples. Nat Neurosci 10:1241–1242. doi:10.1038/nn1961

Dragoi G, Buzsáki G. 2006. Temporal encoding of place sequences by hippocampal cell assemblies. Neuron 50:145–157. doi:10.1016/j.neuron.2006.02.023

Dragoi G, Tonegawa S. 2011. Preplay of future place cell sequences by hippocampal cellular assemblies. Nature 469:397–401. doi:10.1038/nature09633

Drieu C, Zugaro M. 2019. Hippocampal Sequences During Exploration: Mechanisms and Functions. Front Cell Neurosci 13:232. doi:10.3389/fncel.2019.00232

Ecker A, Bagi B, Vértes E, Steinbach-Németh O, Karlócai MR, Papp OI, Miklós I, Hájos N, Freund TF, Gulyás AI, Káli S. 2022. Hippocampal sharp wave-ripples and the associated sequence replay emerge from structured synaptic interactions in a network model of area CA3. Elife11. doi:10.7554/eLife.71850

Enel P, Procyk E, Quilodran R, Dominey PF. 2016. Reservoir Computing Properties of Neural Dynamics in Prefrontal Cortex. PLoS Comput Biol 12:e1004967. doi:10.1371/journal.pcbi.1004967

Feng T, Silva D, Foster DJ. 2015. Dissociation between the experience-dependent development of hippocampal theta sequences and single-trial phase precession. J Neurosci 35:4890–4902. doi:10.1523/JNEUROSCI.2614-14.2015

Fernández-Ruiz A, Oliva A, Nagy GA, Maurer AP, Berényi A, Buzsáki G. 2017. Entorhinal-CA3 Dual-Input Control of Spike Timing in the Hippocampus by Theta-Gamma Coupling. Neuron 93:1213–1226.e5. doi:10.1016/j.neuron.2017.02.017

Foster DJ, Wilson MA. 2007. Hippocampal theta sequences. Hippocampus 17:1093–1099. doi:10.1002/hipo.20345

Foster DJ, Wilson MA. 2006. Reverse replay of behavioural sequences in hippocampal place cells during the awake state. Nature 440:680–683. doi:10.1038/nature04587

Freund TF, Antal M. 1988. GABA-containing neurons in the septum control inhibitory interneurons in the hippocampus. Nature 336:170–173. doi:10.1038/336170a0

Geisler C, Diba K, Pastalkova E, Mizuseki K, Royer S, Buzsáki G. 2010. Temporal delays among place cells determine the frequency of population theta oscillations in the hippocampus. Proc Natl Acad Sci U S A 107:7957–7962. doi:10.1073/pnas.0912478107

Gordon B. 1988. Preserved learning of novel information in amnesia: evidence for multiple memory systems. Brain Cogn 7:257–282. doi:10.1016/0278-2626(88)90002-4

Guan H, Middleton SJ, Inoue T, McHugh TJ. 2021. Lateralization of CA1 assemblies in the absence of CA3 input. Nat Commun 12:6114. doi:10.1038/s41467-021-26389-3

Guzman SJ, Schlögl A, Frotscher M, Jonas P. 2016. Synaptic mechanisms of pattern completion in the hippocampal CA3 network. Science 353:1117–1123. doi:10.1126/science.aaf1836

Hafting T, Fyhn M, Bonnevie T, Moser M-B, Moser EI. 2008. Hippocampus-independent phase precession in entorhinal grid cells. Nature 453:1248–1252. doi:10.1038/nature06957

Haga T, Fukai T. 2018. Recurrent network model for learning goal-directed sequences through reverse replay. Elife7. doi:10.7554/eLife.34171

Hájos N, Karlócai MR, Németh B, Ulbert I, Monyer H, Szabó G, Erdélyi F, Freund TF, Gulyás AI. 2013. Input-output features of anatomically identified CA3 neurons during hippocampal sharp wave/ripple oscillation in vitro. J Neurosci 33:11677–11691. doi:10.1523/JNEUROSCI.5729-12.2013

Harris KD, Henze DA, Hirase H, Leinekugel X, Dragoi G, Czurkó A, Buzsáki G. 2002. Spike train dynamics predicts theta-related phase precession in hippocampal pyramidal cells. Nature 417:738–741. doi:10.1038/nature00808

Henze DA, Urban NN, Barrionuevo G. 2000. The multifarious hippocampal mossy fiber pathway: a review. Neuroscience 98:407–427. doi:10.1016/s0306-4522(00)00146-9

Henze DA, Wittner L, Buzsáki G. 2002. Single granule cells reliably discharge targets in the hippocampal CA3 network in vivo. Nat Neurosci 5:790–795. doi:10.1038/nn887

Hwaun E, Colgin LL. 2019. CA3 place cells that represent a novel waking experience are preferentially reactivated during sharp wave-ripples in subsequent sleep. Hippocampus 29:921–938. doi:10.1002/hipo.23090

Izhikevich EM. 2010. Dynamical Systems in Neuroscience: The Geometry of Excitability and Bursting. The MIT Press.

Jaeger H, Haas H. 2004. Harnessing nonlinearity: predicting chaotic systems and saving energy in wireless communication. Science 304:78–80. doi:10.1126/science.1091277

Jahnke S, Timme M, Memmesheimer R-M. 2015. A Unified Dynamic Model for Learning, Replay, and Sharp-Wave/Ripples. J Neurosci 35:16236–16258. doi:10.1523/JNEUROSCI.3977-14.2015

Jensen O, Idiart MA, Lisman JE. 1996. Physiologically realistic formation of autoassociative memory in networks with theta/gamma oscillations: role of fast NMDA channels. Learn Mem 3:243–256. doi:10.1101/lm.3.2-3.243

Jensen O, Lisman JE. 1996. Theta/gamma networks with slow NMDA channels learn sequences and encode episodic memory: role of NMDA channels in recall. Learn Mem 3:264–278. doi:10.1101/lm.3.2-3.264

Johnson A, Redish AD. 2007. Neural ensembles in CA3 transiently encode paths forward of the animal at a decision point. J Neurosci 27:12176–12189. doi:10.1523/JNEUROSCI.3761-07.2007

Kamondi A, Acsády L, Wang XJ, Buzsáki G. 1998. Theta oscillations in somata and dendrites of hippocampal pyramidal cells in vivo: activity-dependent phase-precession of action potentials. Hippocampus 8:244–261. doi:10.1002/(SICI)1098-1063(1998)8:3<244::AID-HIPO7>3.0.CO;2-J

Karlsson MP, Frank LM. 2009. Awake replay of remote experiences in the hippocampus. Nat Neurosci 12:913–918. doi:10.1038/nn.2344

Kim S, Jung D, Royer S. 2020. Place cell maps slowly develop via competitive learning and conjunctive coding in the dentate gyrus. Nat Commun 11:4550. doi:10.1038/s41467-020-18351-6

King C, Recce M, O’Keefe J. 1998. The rhythmicity of cells of the medial septum/diagonal band of Broca in the awake freely moving rat: relationships with behaviour and hippocampal theta. Eur J Neurosci 10:464–477. doi:10.1046/j.1460-9568.1998.00026.x

Laje R, Buonomano DV. 2013. Robust timing and motor patterns by taming chaos in recurrent neural networks. Nat Neurosci 16:925–933. doi:10.1038/nn.3405

London M, Häusser M. 2005. Dendritic computation. Annu Rev Neurosci 28:503–532. doi:10.1146/annurev.neuro.28.061604.135703

Mehta MR, Lee AK, Wilson MA. 2002. Role of experience and oscillations in transforming a rate code into a temporal code. Nature 417:741–746. doi:10.1038/nature00807

Mehta MR, Quirk MC, Wilson MA. 2000. Experience-dependent asymmetric shape of hippocampal receptive fields. Neuron 25:707–715. doi:10.1016/s0896-6273(00)81072-7

Middleton SJ, McHugh TJ. 2016. Silencing CA3 disrupts temporal coding in the CA1 ensemble. Nat Neurosci 19:945–951. doi:10.1038/nn.4311

Mishra RK, Kim S, Guzman SJ, Jonas P. 2016. Symmetric spike timing-dependent plasticity at CA3– CA3 synapses optimizes storage and recall in autoassociative networks. Nat Commun 7:11552. doi:10.1038/ncomms11552

Miyawaki H, Diba K. 2016. Regulation of Hippocampal Firing by Network Oscillations during Sleep. Curr Biol 26:893–902. doi:10.1016/j.cub.2016.02.024

Mizuseki K, Buzsáki G. 2013. Preconfigured, skewed distribution of firing rates in the hippocampus and entorhinal cortex. Cell Rep 4:1010–1021. doi:10.1016/j.celrep.2013.07.039

Mizuseki K, Royer S, Diba K, Buzsáki G. 2012. Activity dynamics and behavioral correlates of CA3 and CA1 hippocampal pyramidal neurons. Hippocampus 22:1659–1680. doi:10.1002/hipo.22002

Nicola W, Clopath C. 2019. A diversity of interneurons and Hebbian plasticity facilitate rapid compressible learning in the hippocampus. Nat Neurosci 22:1168–1181. doi:10.1038/s41593-019-0415-2

O’Keefe J, Dostrovsky J. 1971. The hippocampus as a spatial map. Preliminary evidence from unit activity in the freely-moving rat. Brain Res 34:171–175. doi:10.1016/0006-8993(71)90358-1

O’keefe J, Nadel L. 1978. The hippocampus as a cognitive map. Oxford university press.

O’Keefe J, Recce ML. 1993. Phase relationship between hippocampal place units and the EEG theta rhythm. Hippocampus 3:317–330. doi:10.1002/hipo.450030307

Oliva A, Fernández-Ruiz A, Buzsáki G, Berényi A. 2016. Spatial coding and physiological properties of hippocampal neurons in the Cornu Ammonis subregions. Hippocampus 26:1593–1607. doi:10.1002/hipo.22659

O’Neill J, Senior TJ, Allen K, Huxter JR, Csicsvari J. 2008. Reactivation of experience-dependent cell assembly patterns in the hippocampus. Nat Neurosci 11:209–215. doi:10.1038/nn2037

Petsche H, Stumpf C, Gogolak G. 1962. The significance of the rabbit’s septum as a relay station between the midbrain and the hippocampus. I. The control of hippocampus arousal activity by the septum cells. Electroencephalogr Clin Neurophysiol 14:202–211. doi:10.1016/0013-4694(62)90030-5

Rebola N, Carta M, Mulle C. 2017. Operation and plasticity of hippocampal CA3 circuits: implications for memory encoding. Nat Rev Neurosci 18:208–220. doi:10.1038/nrn.2017.10

Reifenstein ET, Bin Khalid I, Kempter R. 2021. Synaptic learning rules for sequence learning. Elife10. doi:10.7554/eLife.67171

Rogerson T, Cai DJ, Frank A, Sano Y, Shobe J, Lopez-Aranda MF, Silva AJ. 2014. Synaptic tagging during memory allocation. Nat Rev Neurosci 15:157–169. doi:10.1038/nrn3667

Romani S, Tsodyks M. 2015. Short-term plasticity based network model of place cells dynamics. Hippocampus 25:94–105. doi:10.1002/hipo.22355

Schlesiger MI, Cannova CC, Boublil BL, Hales JB, Mankin EA, Brandon MP, Leutgeb JK, Leibold C, Leutgeb S. 2015. The medial entorhinal cortex is necessary for temporal organization of hippocampal neuronal activity. Nat Neurosci 18:1123–1132. doi:10.1038/nn.4056

Schlingloff D, Káli S, Freund TF, Hájos N, Gulyás AI. 2014. Mechanisms of sharp wave initiation and ripple generation. J Neurosci 34:11385–11398. doi:10.1523/JNEUROSCI.0867-14.2014

Scoville WB, Milner B. 1957. Loss of recent memory after bilateral hippocampal lesions. J Neurol Neurosurg Psychiatry 20:11–21. doi:10.1136/jnnp.20.1.11

Sharif F, Tayebi B, Buzsáki G, Royer S, Fernandez-Ruiz A. 2021. Subcircuits of Deep and Superficial CA1 Place Cells Support Efficient Spatial Coding across Heterogeneous Environments. Neuron 109:363–376.e6. doi:10.1016/j.neuron.2020.10.034

Silva D, Feng T, Foster DJ. 2015. Trajectory events across hippocampal place cells require previous experience. Nat Neurosci 18:1772–1779. doi:10.1038/nn.4151

Skaggs WE, McNaughton BL, Wilson MA, Barnes CA. 1996. Theta phase precession in hippocampal neuronal populations and the compression of temporal sequences. Hippocampus 6:149–172. doi:10.1002/(SICI)1098-1063(1996)6:2<149::AID-HIPO6>3.0.CO;2-K

Stimberg M, Brette R, Goodman DF. 2019. Brian 2, an intuitive and efficient neural simulator. Elife8. doi:10.7554/eLife.47314

Sussillo D, Abbott LF. 2009. Generating coherent patterns of activity from chaotic neural networks. Neuron 63:544–557. doi:10.1016/j.neuron.2009.07.018

Suzuki SS, Smith GK. 1988. Spontaneous EEG spikes in the normal hippocampus. III. Relations to evoked potentials. Electroencephalogr Clin Neurophysiol 69:541–549. doi:10.1016/0013-4694(88)90166-6

Tiesinga P, Sejnowski TJ. 2009. Cortical enlightenment: are attentional gamma oscillations driven by ING or PING? Neuron 63:727–732. doi:10.1016/j.neuron.2009.09.009

Tolman EC. 1948. Cognitive maps in rats and men. Psychol Rev 55:189–208. doi:10.1037/h0061626

Tsodyks MV, Skaggs WE, Sejnowski TJ, McNaughton BL. 1996. Population dynamics and theta rhythm phase precession of hippocampal place cell firing: a spiking neuron model. Hippocampus 6:271–280. doi:10.1002/(SICI)1098-1063(1996)6:3<271::AID-HIPO5>3.0.CO;2-Q

Wang JX, Kurth-Nelson Z, Kumaran D, Tirumala D, Soyer H, Leibo JZ, Hassabis D, Botvinick M. 2018. Prefrontal cortex as a meta-reinforcement learning system. Nat Neurosci 21:860–868. doi:10.1038/s41593-018-0147-8

Weber SN, Sprekeler H. 2018. Learning place cells, grid cells and invariances with excitatory and inhibitory plasticity. Elife 7. doi:10.7554/eLife.34560

Yamamoto J, Tonegawa S. 2017. Direct Medial Entorhinal Cortex Input to Hippocampal CA1 Is Crucial for Extended Quiet Awake Replay. Neuron 96:217–227.e4. doi:10.1016/j.neuron.2017.09.017

Yassa MA, Stark CEL. 2011. Pattern separation in the hippocampus. Trends Neurosci 34:515–525. doi:10.1016/j.tins.2011.06.006

Zutshi I, Brandon MP, Fu ML, Donegan ML, Leutgeb JK, Leutgeb S. 2018. Hippocampal Neural Circuits Respond to Optogenetic Pacing of Theta Frequencies by Generating Accelerated Oscillation Frequencies. Curr Biol 28:1179–1188.e3. doi:10.1016/j.cub.2018.02.061

